# Endocytic Clathrin Coats Develop Curvature at Early Stages of Their Formation

**DOI:** 10.1101/715219

**Authors:** Nathan M. Willy, Joshua P. Ferguson, Salih Silahli, Cemal Cakez, Farah Hasan, Yan Chen, Min Wu, Henry C. Chang, Alex Travesset, Siyu Li, Roya Zandi, Dong Li, Eric Betzig, Emanuele Cocucci, Comert Kural

**Affiliations:** Department of Physics, Ohio State University, Columbus, OH 43210; Department of Nuclear Engineering, University of New Mexico, Albuquerque, NM 87131; Centre for Bioimaging Sciences, Department of Biological Sciences, National University of Singapore, Singapore; Mechanobiology Institute, National University of Singapore, Singapore; Department of Biological Sciences, Purdue University, West Lafayette, IN 47907; Department of Physics and Astronomy, Iowa State University, Ames, IA 50011; Ames Laboratory, Iowa State University, Ames, IA 50011; Department of Physics and Astronomy, University of California, Riverside, CA 92521; National Laboratory of Biomacromolecules, CAS Center for Excellence in Biomacromolecules, Institute of Biophysics, Chinese Academy of Sciences, Beijing, China 100101; College of Life Sciences, University of Chinese Academy of Sciences, Beijing, China 100049; Departments of Physics and Molecular and Cell Biology, University of California, Berkeley, CA, 94720; Janelia Research Campus, Howard Hughes Medical Institute, Ashburn, VA, 20147; College of Pharmacy, Ohio State University, Columbus, OH 43210; Interdisciplinary Biophysics Graduate Program, Ohio State University, Columbus, OH 43210

## Abstract

Sculpting a flat patch of membrane into an endocytic vesicle requires curvature generation on the cell surface, which is the primary function of endocytic protein complexes. The mechanism through which membrane curvature is imposed during formation of clathrin-coated vesicles is an ongoing controversy. Using super-resolved live cell fluorescence imaging, we demonstrate that curvature generation by clathrin-coated pits can be detected in real time within cultured cells and tissues of developing metazoan organisms. We found that the footprint of clathrin coats increase monotonically during formation of curved pits at different levels of plasma membrane tension. Our findings are only compatible with models that predict curvature generation at early stages of endocytic clathrin-coated pit formation. Therefore, clathrin-coated vesicle formation does not necessitate a dynamically unstable clathrin lattice that would allow an abrupt flat-to-curved transition.

**Summary:** Endocytic clathrin coats acquire curvature without a flat-to-curved transition that requires an extensive reorganization of the clathrin lattice.

---

Clathrin-mediated endocytosis is the most extensively studied internalization mechanism of membrane lipids and proteins from the cell surface ^1,2^. Over the past decades, a multitude of biophysical and biochemical methodologies have been employed to elucidate structural and dynamic properties of endocytic clathrin coats ^3^. However, fundamental aspects of clathrin-mediated endocytosis remain controversial due to the lack of experimental approaches that allow correlation of ultra-structural and dynamic properties of assembling clathrin coats. Clathrin triskelions can assemble into polyhedral cages and hexagonal lattices in seemingly infinite numbers of geometries upon their recruitment to the plasma membrane by a range of adaptor proteins including AP2 ^4,5^. Based on their structural and dynamic properties, clathrin coats are categorized into pits and plaques ^6,7^. Clathrin-coated pits are highly curved structures with hexagonal and pentagonal faces. They are internalized in the form of coated vesicles within few minutes of their initiation at the plasma membrane ^8–10^ (Fig. 1A). Due to their small size (100-200 nm in diameter), clathrin pits appear as diffraction-limited spots under conventional fluorescence imaging^11^. Plaques are large and flat hexagonal lattices of clathrin that are longer-lived than pits ^7,12,13^.

**Figure 1.**
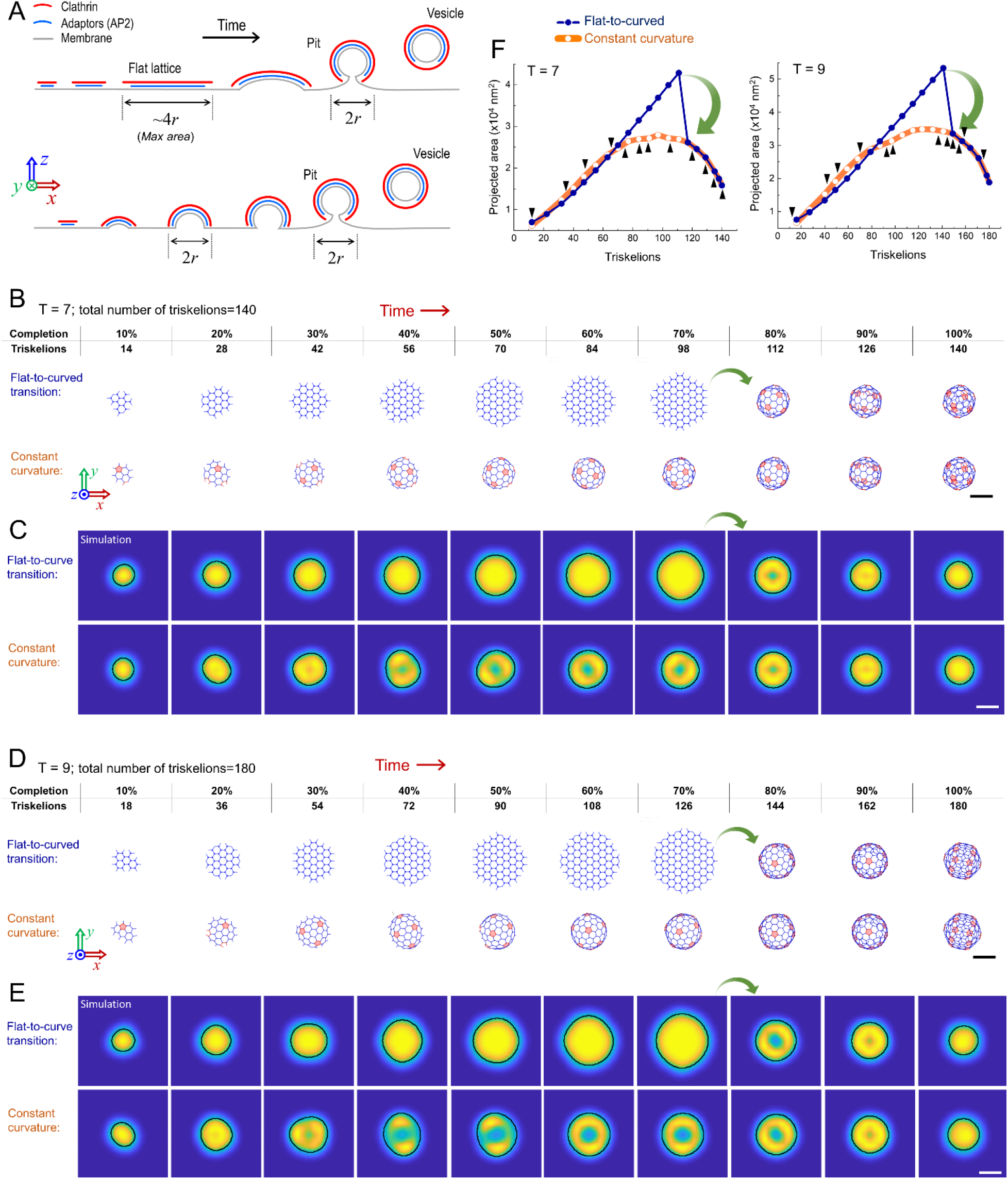
Flat-to-curved transition necessitates a substantial change in the projected area of clathrin coats. **A.** Schematics depicting the cross-sections of clathrin coats as suggested by prevalent curvature generation models. According to the flat-to-curved transition models (upper), the clathrin coat initially grows into a flat hexagonal lattice constituting 70-100% of the final surface area of the coated vesicle (4π*r*^2^; *r* is the radius of the pit/vesicle). A disc-shaped, flat clathrin lattice is anticipated to reach a diameter of 3*r*-4*r* prior to the transition into a pit ^4,18,19^. Upon this transition, the projected coat area reduces to π*r*^2^. The constant curvature model (lower) predicts a gradual increase in the projected coat area until the diameter reaches 2*r*, which is also the diameter of the clathrin pit/vesicle. **B.** 140 triskelions are assembled into a T=7 polyhedron through a flat-to-curved transition (marked by green arrow) taking place at 70% of growth (upper panel) and as predicted by the constant curvature model (lower panel; movie S1). Pentagonal faces are highlighted in pink. **C.** Snapshots of high-NA TIRF-SIM simulations correspond to the distinct completion levels of T=7 clathrin coats in (B). Detected boundaries are shown by the black demarcations for both flat-to-curved transition and constant curvature models. **D.** 180 triskelions are assembled into a T=9 polyhedron through a flat-to-curved transition (marked by green arrow) taking place at 70% of growth (upper panel) and as predicted by the constant curvature model (lower panel; movie S2). Pentagonal faces are highlighted in pink. **E.** Snapshots of high-NA TIRF-SIM simulations correspond to the distinct completion levels of T=9 clathrin coats in (D). Detected boundaries are shown by the black demarcations for both flat-to-curved transition and constant curvature models. **F.** Detected area of the structures obtained from high-NA TIRF-SIM simulations (C & E) are plotted with respect to number of triskelions for both models. Green arrows mark the significant reduction in the detected area upon the proposed flat-to-curved transition at 70% of coat growth. Black arrowheads mark completion of pentagonal faces predicted in the constant curvature model. The difference between the two models is evident independent of the distance between the coverslip and the coat (Fig. S5). Scale bars, 100 nm.

Formation of endocytic vesicles requires curvature generation during the lifespan of clathrin coats. However, how and at what point of its growth a clathrin coat develops curvature has been a matter of debate for almost four decades ^14–16^. Currently, there are two competing models as to how clathrin and its accessory proteins generate curvature that leads to formation of clathrin-coated pits. It was originally proposed that clathrin initially grows into a flat hexagonal array on the plasma membrane prior to transitioning into a curved coat ^4,17^ (Fig. 1A, top). This flat growth is postulated to reach 70-100% of the final surface area before remodeling into a clathrin-coated pit through transformation of the hexagonal clathrin lattice into a curved polyhedron with pentagonal and hexagonal faces ^4,18,19^. Eventually, 12 pentagonal faces must be introduced to generate a closed clathrin cage ^20,21^ (Fig. 1B & D). This model was rejected by others because insertion of even a single pentagonal face into a hexagonal lattice requires a substantial structural rearrangement (see Figure 13 of^4^), which is energetically unfavorable ^14,15,22^. As an alternative, it was suggested that clathrin-coated pits form gradually without a significant structural rearrangement. According to this model, as clathrin triskelions are integrated into the coat, they organize into pentagonal as well as hexagonal faces to generate a constant amount of curvature (Fig. 1A, bottom). The debate regarding curvature formation by clathrin pits has been recently rekindled by studies employing correlative fluorescence and electron microscopy of clathrin-coated structures ^16,18,19,23^. However, in these assays, curvature formation by individual clathrin coats could not be monitored directly due to absence of temporal dimension in electron microscopy acquisitions.

Here we used super-resolved fluorescence microscopy to image endocytic clathrin coat dynamics in live cells and tissues. Enhanced resolution in both spatial and temporal domains allowed us to monitor curvature generation by clathrin-coated pits in real time. Detailed analyses of the footprint and the fluorescence intensity of the assembling clathrin coats have revealed that these structures do not undergo major structural rearrangements proposed by the flat-to-curved transition models even under increased membrane tension. We also used single-molecule detection of clathrin triskelions in cell-free reconstitution systems to show that clathrin coats do not exhibit global dynamic instability that would allow flat-to-curved transitions. Overall, our results show that curvature is generated at very early stages of clathrin coat formation, in good agreement with the constant curvature model.

## Results

### Flat-to-curved transition necessitates a significant decrease in the footprint of clathrin coats

One of the major observables that distinguish the flat-to-curved transition and constant curvature models is the change in the projected area (i.e., footprint) of the clathrin coat as it gains curvature: While the constant curvature model predicts a steady rise in the projected area of the coat on the imaging plane, the flat-to-curved transition requires an abrupt three- to four-fold reduction in the same measure, corresponding to transition at 70% (see Figure 1 of^19^) or 100% of coat growth (see Figure 6 of^4^) as proposed in previous studies. To demonstrate this, we adapted a previously described self-assembly model to simulate growth of clathrin polyhedra with triangulation (T) numbers of 7 and 9 (Figure 1B-F; Movies S1 & S2) ^24^. We chose to work with these specific geometries as their dimensions are in good agreement with clathin pits reported in previous electron microscopy studies ^4,18,19,25,26^. In Figure 1B, 140 triskelions are assembled into a T=7 polyhedron (60 hexagons and 12 pentagons; ~137 nm diameter and ~14,700 nm^2^ final projected coat area) via a flat-to-curved transition taking place at 70% of completion (upper panel)^19^, and as predicted by the constant curvature model (lower panel). The size of the simulated T=7 polyhedron is an accurate representation of clathrin-coated pits detected in BSC-1 cells^19^, human embryonic stem cells (hESCs) and derived neuronal progenitor cells (NPCs)^27^. Figure 1D shows assembly of a larger polyhedron (T=9) comprising 180 triskelions (80 hexagons and 12 pentagons; ~156 nm diameter and ~19,100 nm^2^ final projected coat area) using both models. The large footprint of the T=9 polyhedon is a good representation of clathrin-coated pits detected in and fibroblasts differentiated from hESCs^27^. In both scenarios, flat-to-curved transition (marked with green arrows) results in significant decrease in the footprint of the clathrin coat (Figure 1B & D). Note that the change in the footprint would be even more pronounced if the transition took place after the full assembly (100% completion) of the coat, as proposed earlier (i.e., constant area model) ^18^.

Electron microscopy provides high-resolution snapshots of clathrin-coated structures with different curvature levels ^4,18,19,26^. However, due to the lack of temporal dimension, electron micrographs fail to provide direct evidence for the mechanism of clathrin-driven curvature generation. Conventional live cell fluorescence imaging allowed researchers to elucidate the formation and dissolution dynamics of endocytic clathrin coats in living cells ^9,11,28^. In these studies structural properties of clathrin coats are obscured by diffraction, which limits spatial resolution. Nevertheless, diffraction limit in fluorescence imaging can be circumvented by recently developed superresolution microscopy techniques, such as Stochastic Optical Reconstruction Microscopy (STORM) and Structured Illumination Microscopy (SIM). In these assays, curved clathrin coats (i.e. pits) appear as rings, which correspond to two-dimensional projection of spheroids ^29–32^. Using the triskelion coordinates in our self-assembly models, we simulated high-NA TIRF-SIM (structured illumination microscopy in the total internal reflection fluorescence mode) images of T=7 and T=9 polyhedra at different completion levels. As expected, ring patterns developed in the TIRF-SIM simulations upon formation of clathrin pits. We used an edge detection algorithm to find the boundaries of the simulated TIRF-SIM images (Figs. 1C & E). Since our simulations incorporate fluorescence diffraction, the area enclosed within the detected boundaries of the simulated TIRF-SIM images appear larger than the footprint of the corresponding polyhedron. Due to the substantial change in the coat geometry, the projected area of the TIRF-SIM images decreased significantly upon the simulated flat-to-curved transitions taking place at 70% of coat completion^19^ (Figs. 1F & S5).

### The projected area of clathrin coats increase monotonically until formation of pits as predicted by the constant curvature model

We employed TIRF-SIM to image the formation of clathrin-coated pits in cultured cells (in vitro) and tissues of developing *Drosophila* melanogaster embryos (in vivo) with high spatial and temporal resolution and monitor curvature generation by endocytic clathrin coats in real time ^31^ (Movie S3). In these assays, the area of clathrin coat images increased monotonically until the ring pattern arises, which marks formation of clathrin-coated pits. This is followed by a relatively fast disappearance of fluorescence due to uncoating (Fig. 2). To perform a bulk analysis, we developed a custom tracking software for manual selection of individual clathrin-coated pit traces that appear independent of other structures from their formation to dissolution, and contain a single phase of fluorescence intensity growth followed by a decline. We created the time average of clathrin-coated pit traces extracted from BSC-1 and COS-7 cells imaged with high-NA TIRF-SIM, and calculated the temporal evolution of the detected area (Movies S4 & S5; Figs. 3A) and the size of the ring pattern (i.e., peak-to-peak separation of the radial average; Fig. 3B). Contrary to the predictions of the flat-to-curved transition models, we detected no decrease in the footprint of the coat prior to appearance of the ring pattern, which marks development of a clathrin-coated pit. Instead, we found that the ring pattern is observable as the area makes a plateau and reaches the maximum (Fig. 3A, B), as predicted by the constant curvature model (Fig. 1F). Maximum average area detected from BSC-1 and COS-7 clathrin pit images correlate with the TIRF-SIM simulations developed for T=7 and T=9 polyhedra, respectively (Figs. 1F & 3A). These results signify the accuracy of our measurements given that the size of the T=7 polyhedron is in perfect agreement with the average clathrin pit dimensions reported in previous electron microscopy assays performed on BSC-1 cells ^19^.

**Figure 2.**
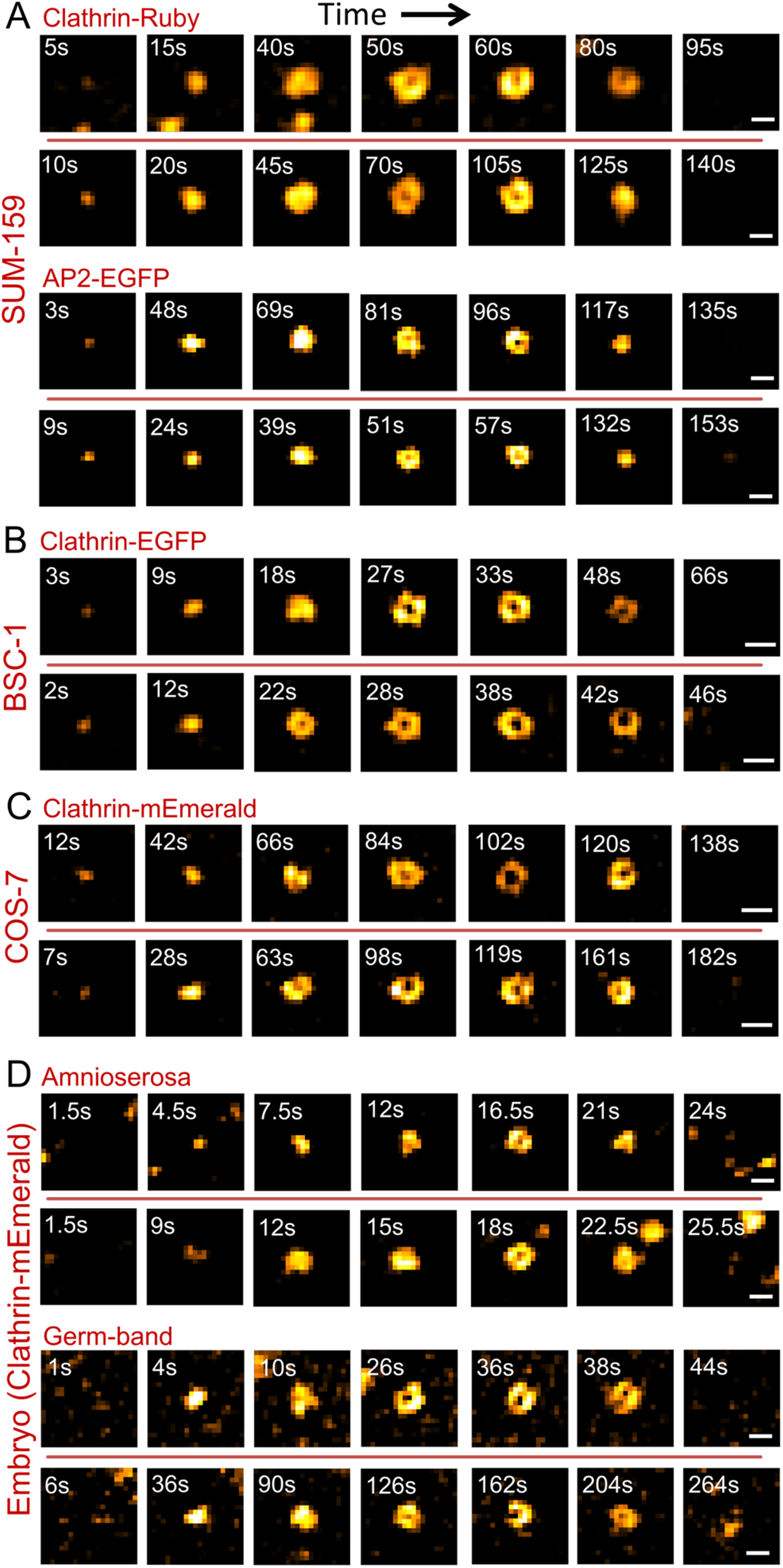
TIRF-SIM allows analysis of curvature generation by clathrin pits with high spatiotemporal resolution in cultured cells and in tissues of Drosophila embryos. Each row shows snapshots from individual clathrin-coated pit traces. Formation of pits (marked by a ring pattern) can be observed under various *in vitro* and *in vivo* conditions including SUM-159 cells expressing clathrin-Ruby and AP2-EGFP as endocytic markers (A), BSC-1 cells expressing clathrin-EGFP (B), COS-7 cells expressing clathrin-mEmerald (C), and amnioresora and germ-band tissues of Drosophila embryos expressing clathrin-mEmerald (D). Two traces are shown for each condition. Scale bars, 200 nm.

**Figure 3.**
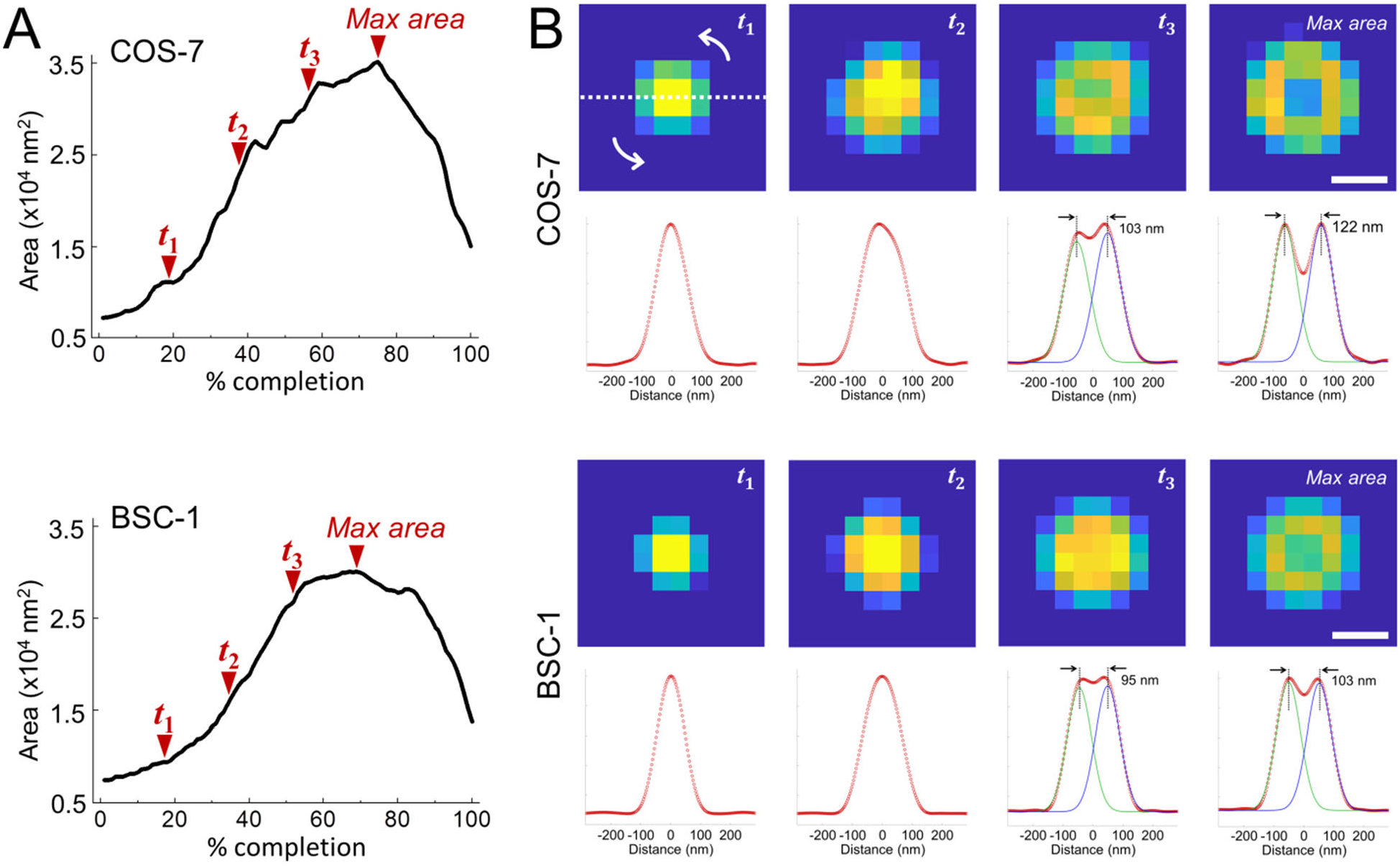
Clathrin coats reach maximum projected area by formation of pits. **A.** Average clathrin-coated pit formation and dissolution images using 34 and 81 individual traces are obtained from COS-7 cells expressing clathrin-mEmerald and BSC-1 cells expressing clathrin-EGFP, respectively (Movies S4 & S5). In both cases, the area of the average image increases monotonically until a plateau is reached, which is followed by a relatively sharp decrease. **B.** For both cell types, average images of clathrin coats are shown for four equally spaced % completion points, last of which is the maximum area point (denoted as *t*_1_, *t*_2_, *t*_3_ and *max area* in A). Note that the ring pattern becomes apparent as the area converges to the maximum. Below each image is the radial average (red circles), which is obtained along a cross-section rotated 180° around the center. We used two Gaussian fits to the radial average (green and blue) to calculate the peak-to-peak separation as an estimate of the ring size. Scale bars, 90 nm.

As an alternative to manual selection, we developed an algorithm for automated detection of individual clathrin-coated pit traces that appear independent of other clathrin structures in TIRF-SIM acquisitions. To ensure detection of clathrin-coated pit formation, these automatically selected traces had to have the ring pattern appear in at least two consecutive frames. The first appearance of the ring pattern in the trace is marked as the “ring frame”, which is used for synchronization of traces and sorting them into three groups based on area (i.e., small, medium and large). Figure 4A shows the average of synchronized traces (extending to 5 frames before and after the ring frame) detected in BSC-1 cells imaged at 0.5 frames/second. In good agreement with the manually selected traces, we found that the footprint of the structures increases and reaches a plateau with the appearance of the ring pattern in all three groups (Figs. 4B, C). The results of the same analysis are shown in Figures S2-S4 for COS-7 cells and BSC-1 cells imaged at different frame rates. Overall, contrary to the predictions of the flat-to-curved transition models, we found that curvature generation is not associated with reduction of the projected area, i.e. the footprint of the coats increases throughout maturation of clathrin-coated pits in accord with the constant curvature model (Fig. 4D).

**Figure 4.**
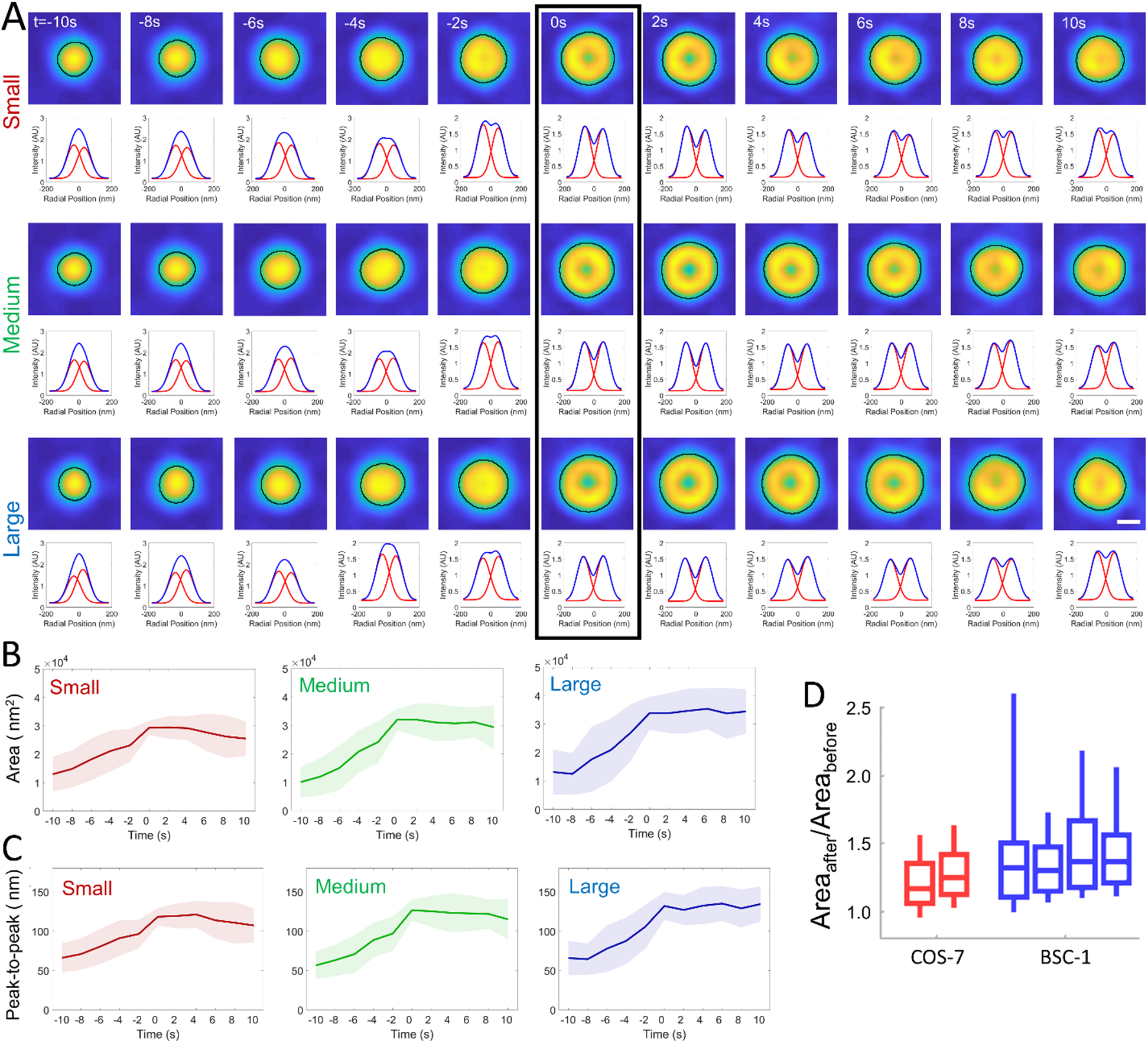
Automated analysis of the projected area during formation of clathrin pits. **A.** 187 traces detected from two BSC-1 cells (imaged at 0.5 frames/second) are grouped and synchronized using the first detected ring pattern (black rectangle). Average images (top) and corresponding radial averages (bottom) are shown for each group (small, medium and large). Detected boundaries are shown by the black demarcations. The temporal evolution of the average area **(B)** and peak-to-peak separation of the Gaussian fits to the radial averages **(C)** are plotted for the three groups. Shaded areas represent the standard deviation. **D.** Boxplots show the ratio of the maximum projected areas detected after and before the ring frame (Area_after_/Area_before_) in clathrin traces automatically detected from two COS-7 (red; *N*_traces_ = 694) and four BSC-1 cells (*N*_traces_ = 397) imaged with high-NA TIRF-SIM. Overwhelming majority of the clathrin structures reach maximum area upon formation of pits, in good agreement with the manual analysis (Figure 3). Boxes extend to the quartiles, with a line at the median. Whiskers extend from the 10th to 90th percentiles. Scale bar, 100 nm.

### The clathrin coat assembly and invagination are synchronous as predicted by the constant curvature model

Transition of a flat clathrin lattice into a curved pit necessitates a significant inward movement that will displace the center of mass of the coat away from the membrane/glass interface (Fig. 5A, top). Therefore, due to the evanescent profile of the high-NA TIR illumination (i.e., 50 nm penetration depth corresponding to excitation NA of 1.55^31^), our simulations predicted a significant (~3-fold) reduction in the integrated fluorescence intensity of the clathrin coat upon flat-to-curved transition (Fig. 5B). The same outcome was observed in recent simulations of Scott et al., where they adopted a longer penetration depth (100 nm) applicable to the lower excitation NA used in their TIR imaging (see Supplementary Figure 1c of ^33^). In contrast, the constant curvature model proposes that the assembly of the clathrin coat is synchronous with its invagination and, therefore, the apex of the coat departs from the glass surface continuously (Fig. 5A, bottom). Correspondingly, clathrin fluorescence is predicted to converge into a plateau as the coat matures into a pit ^7,34^ (Fig. 5B). Note that the striking difference between the two models is obvious regardless of the distance between the clathrin coat and the substrate (Fig. S5). We used automatically selected clathrin coat traces that are synchronized to the ring frame to analyze the temporal evolution of the fluorescence signal. In BSC-1 and COS-7 cells imaged at different frame rates, we detected no reduction in the fluorescence signal upon the appearance of the ring pattern. Instead, the signal increased and converged into a plateau in good agreement with the constant curvature model (Fig. 5C). Similar plateau patterns were reported in earlier studies that employed TIR illumination to characterize fluorescence intensity profiles of clathrin-coated pits^7,9,34^.

**Figure 5.**
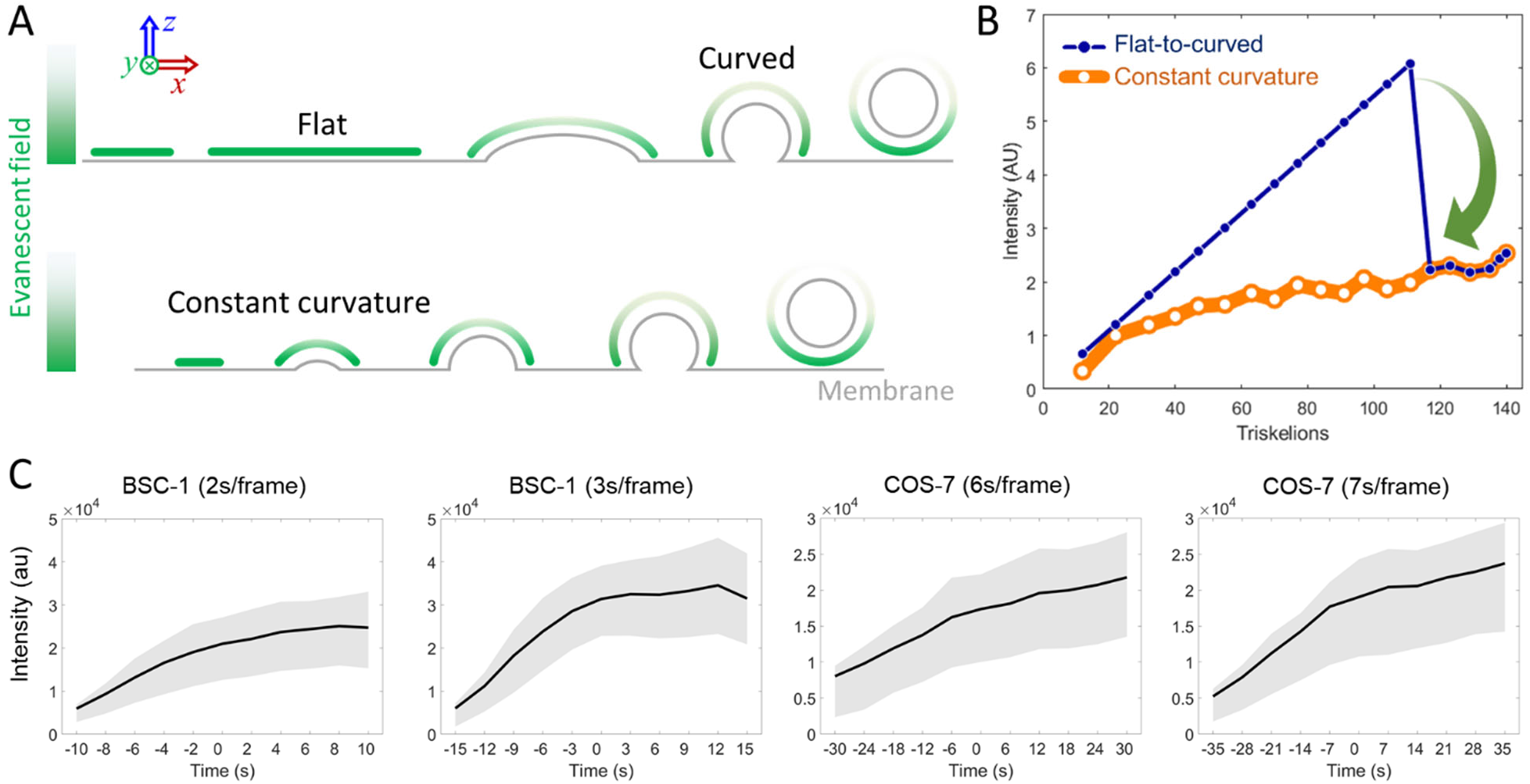
**A.** Schematics depicting illumination of clathrin coats under evanescent field created by the total internal reflection (TIR) of excitation laser. Flat coats are illuminated with higher intensity (green) since they are closer to the glass/sample interface (above). Invagination moves the apex of the coat away from the interface, which is synchronous to coat growth according to the constant curvature model (below). **B.** Simulations show integrated TIRF intensity as a function of coat completion for T=7 polyhedra assembled based on the two models. Constant curvature model predicts a plateau in the clathrin signal (orange), whereas a late stage flat-to-curved transition (at 70% of completion) results in a significant decrease in intensity (blue). Penetration depth for high-NA TIRF illumination is 50 nm. **C.** TIRF intensity traces (average and standard deviation) are plotted for automatically selected and synchronized traces from BSC-1 cells and COS-7 cells imaged at different frame rates. The ring frame corresponds to time = 0s.

**Figure 6.**
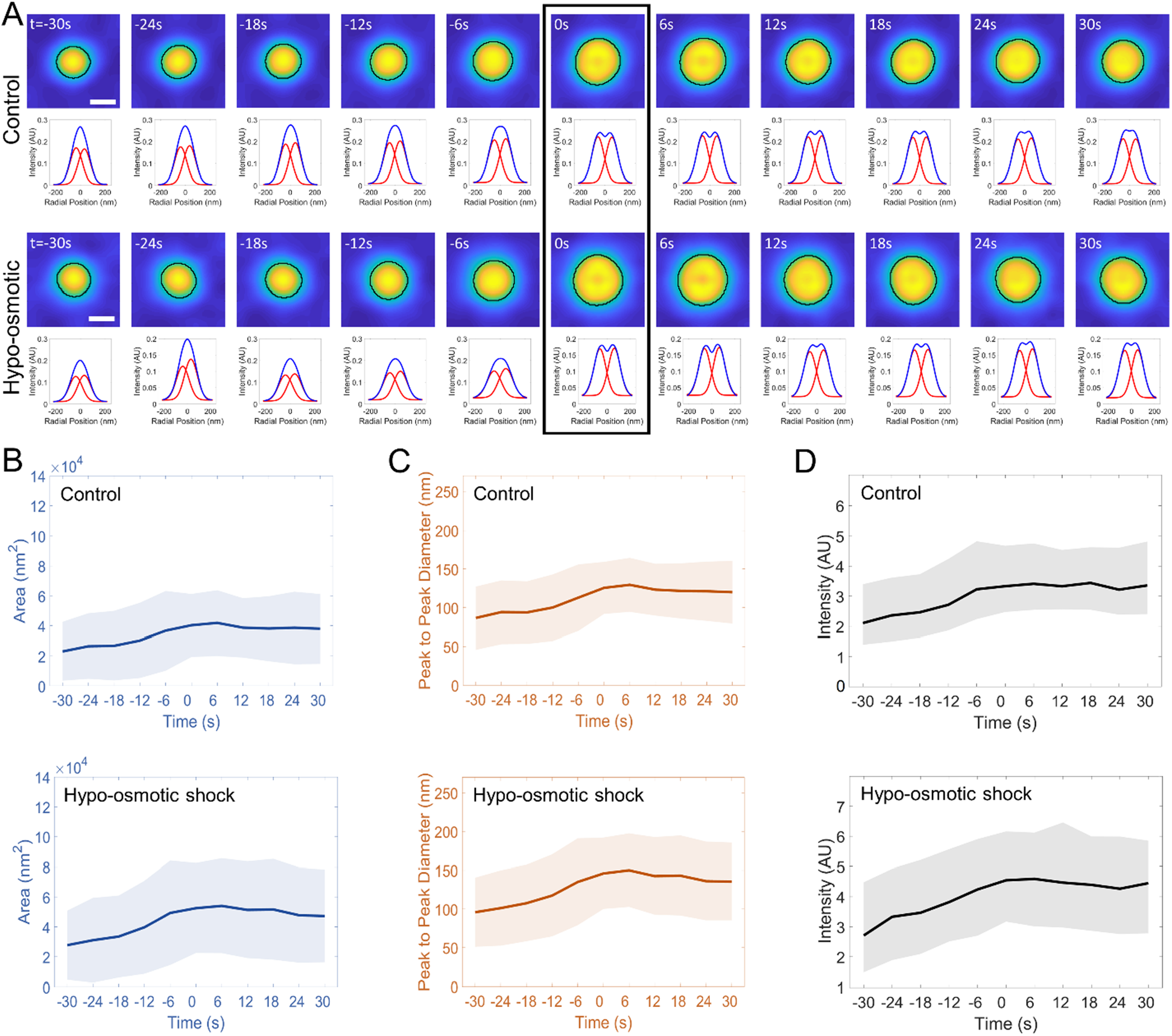
Curvature generation by clathrin coats under high plasma membrane tension. Membrane tension is increased in cells by hypo-osmotic shock. Clathrin traces are automatically selected in SUM-159 cells genome edited to express AP2-EGFP under isotonic (*N*_cells_ = 3; *N*_traces_ = 366) and hypotonic (*N*_cells_ = 9; *N*_traces_ = 396) conditions. **A.** The average of the synchronized traces and corresponding radial averages are shown for both conditions. The arrowhead denotes the ring frame. The temporal evolution of the average area **(B)** and peak-to-peak separation of the Gaussian fits to the radial averages **(C)** are plotted for the two groups. Shaded areas represent the standard deviation. **D.** TIRF intensity traces (average and standard deviation) are plotted for automatically selected and synchronized traces from SUM-159 cells imaged at isotonic and hypotonic conditions. Scale bars, 100 nm.

### Membrane tension does not alter the mechanism of curvature generation

Plasma membrane tension is one of the most important determinants of endocytic clathrin coat structure and dynamics as it increases the energetic cost of membrane deformation ^19,35–38^. Our automated analysis on genome edited SUM-159 cells expressing AP2-EGFP^39^ revealed that average size of clathrin-coated pits increase when plasma membrane tension is elevated by hypo-osmotic shock (Fig. 6A-C), confirming previous reports ^37^. Regardless of tension level, both detected area and fluorescence signal increased monotonically throughout the maturation of clathrin pits and reached a plateau (Fig. 6B- D). These results show that increased plasma membrane tension does not induce a late stage flat-to-curved transition that leads to formation of clathrin-coated pits.

### Dynamic instability is localized to the edge of assembling clathrin coats

Flat-to-curved transition of the clathrin coat requires a major structural reorganization that disrupts the hexagonal arrangement of the flat clathrin lattice to introduce pentagonal faces ^4,14,15^. It was proposed that dynamic instability throughout the entire coat surface allows such transitions through rapid exchange of clathrin triskelions ^18^. Using dual color single-molecule imaging of clathrin dissociations from assembling pits (Fig. 7A), we have recently shown that the dynamic exchange of clathrin serves as a proofreading mechanism for functional cargo incorporation, rather than facilitating curvature transitions^40^. Moreover, the short dwell times of transient clathrin recruitments in these assays suggest that dissociation events are localized to the edge of growing pits^40^. To further demonstrate the difference, for both of these scenarios (i.e., global and edge instability, respectively), we simulated the probability of clathrin dissociations based on the self-assembly model developed for T=7 polyhedron, and compared the results with single-molecule data we obtained from cell-free reconstitution assays. The simulated frequency of dissociations increased steadily until the end of coat completion when we allowed dissociations to take place anywhere on the coat surface (global instability). On the other hand, when dissociations are limited to the edge of the growing coat (edge instability), we found that the frequency reaches a peak halfway through the completion, i.e. when the neck/edge is the widest (Fig. 7B). Our experimental results coincide with the latter scenario; frequency of transient clathrin recruitments increase and decrease with the expansion and constriction of the neck region, respectively (Fig. 7C). We also performed fluorescence recovery after photobleaching (FRAP) assays on clathrin coats that grow synchronously in cell-free reconstitution assays^40^. We found that as the clathrin pits reach the most mature state there is essentially no fluorescence recovery, further evidence against curvature generation models that propose a globally instable clathrin coat (Fig. 7D, E).

**Figure 7.**
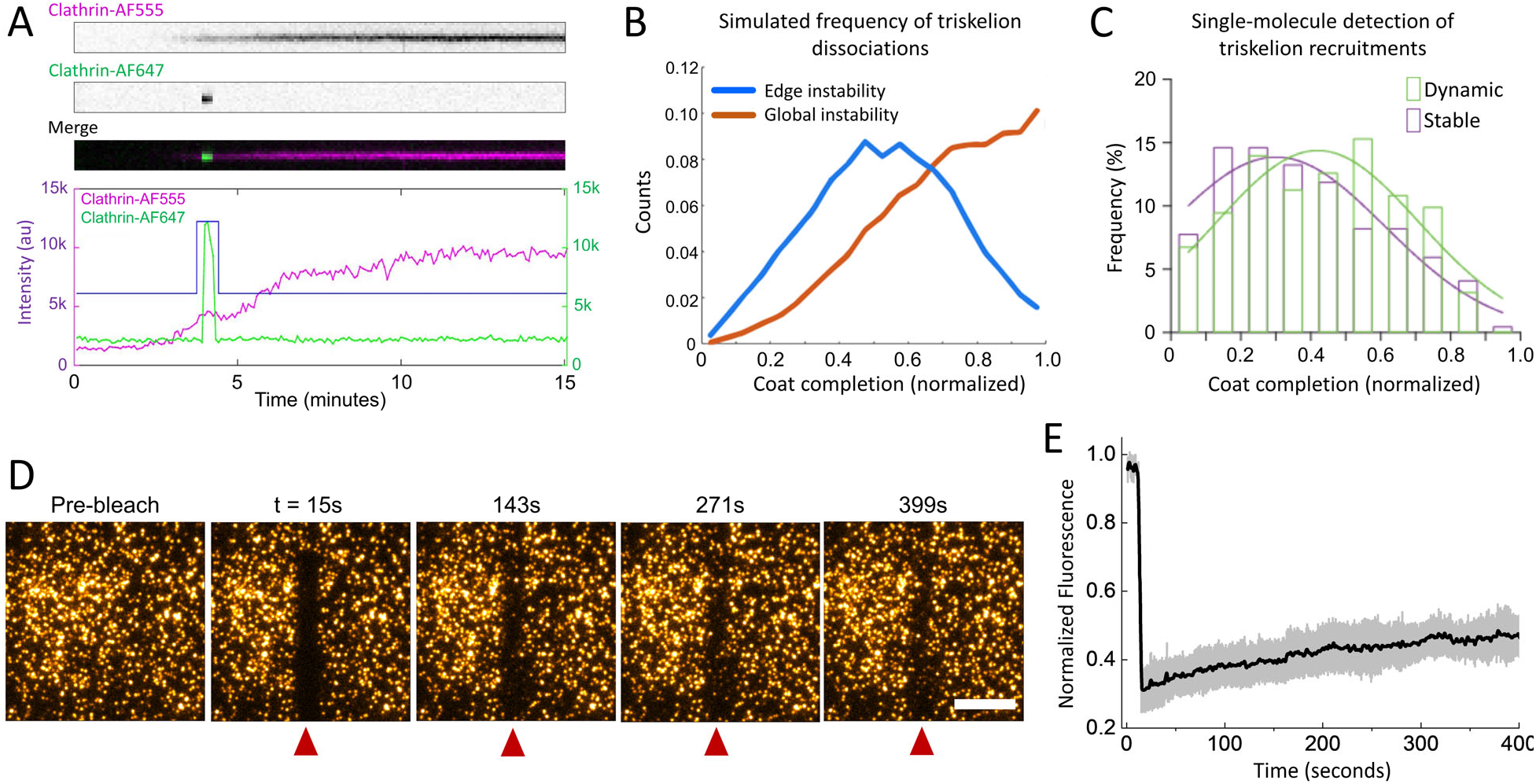
Dynamic instability is limited to the edges of growing clathrin pits. **A.** The three kymographs show dynamic (i.e., transient) recruitment of an Alexa Fluor 647-labeled clathrin triskelion (Clathrin-AF647) to an assembling pit that is visualized by Alexa Fluor 555-labeled clathrin (Clathrin-AF555) signal. Fluorescence intensity plots of the two images (bottom) show assembly of the clathrin pit (purple) and the transient recruitment (green). **B.** Frequency of triskelion exchange is expected to be proportional to the area of a globally instable clathrin coat (red). Edge instability predicts the exchange frequency to rise and fall with the perimeter of the coat edge (blue). **C.** Single molecule detection of triskelion recruitments show that transient (dynamic) events are more frequently observed halfway through the completion of the clathrin coats, i.e. when the edge is the widest. **D.** FRAP assays are performed on clathrin-coated pits that have matured on cell-free reconstitution systems. Arrowheads mark the bleached region at different time points. **E.** Fluorescence recovery curve (normalized; mean ± standard deviation) is obtained from 37 clathrin puncta photobleached after completion of growth. Scale bar, 10 *μ*m.

## Discussion

Here we analyzed clathrin-coated pit formation with high spatial and temporal resolution in a wide range of cell types as well as, for the first time, tissues of developing organisms. Flat-to-curved transition models suggest extensive alterations in coat structure which necessitate substantial changes in footprint and fluorescence intensity prior to formation of clathrin pits. Our findings clearly demonstrate that curvature generation by clathrin-coated pits takes place through a mechanism that is independent of such a transition. Even though our findings are compatible with the constant curvature model, we do not claim that curvature has to be uniform throughout the coat; heterogeneous insertion of the pentagonal faces can result in formation of clathrin coats that do not exhibit icosahedral symmetry^41^. We have previously shown that Hsc70-driven dynamic instability at the edges of assembling clathrin pits serves as a proofreading mechanism for cargo enrichment^40^. Physical properties of plasma membrane and cargo (e.g., membrane tension, cargo size and density) might dictate coat curvature, and therefore, the size of assembling pits through this mechanism^36,42^.

It was previously proposed that a globally unstable clathrin coat would allow a late stage flat-to-curved transition^18^. This assumption was supported by fast fluorescence recovery rates detected after photobleaching of fluorescent clathrin spots in living cells. It is important to note that a majority of the fluorescent spots that appear as individual clathrin coats under diffraction-limited imaging actually contain multiple independent structures that are positioned in close proximity (Fig. S6, Movie S6). Therefore, it is worthwhile to consider that the detected fluorescence recovery may actually correspond to initiation and growth of adjacent clathrin coats that cannot be resolved as separate structures in these assays. Indeed, our FRAP assays performed on cell-free reconstitution systems show that there is no fluorescence recovery as clathrin pits reach the most mature state (Fig. 7D & E).

In a recent study, Scott et al. utilized polarized TIRF to characterize curvature generation by endocytic clathrin coats^33^. Using the ratio of fluorescence signals obtained by parallel (S) and perpendicular (P) excitation of DiI molecules incorporated into the plasma membrane (i.e. P/S), they concluded that a small fraction of clathrin coats undergo a late stage flat-to-curved transition. However, in this population of traces, the fluorescence of the clathrin signal plateaus prior to the detection of a rise in P/S, which is incompatible with the flat-to-curved transition model. Instead, as demonstrated by their own flat-to-curved simulations (supplementary figure 1c in^33^) and ours (Fig. 5B), the invagination of the coat (marked by an increase in P/S) must coincide with almost two-fold reduction in the clathrin intensity instead of a plateau. Since this population of traces have longer lifetimes, we believe that they belong to larger (less curved) clathrin-coated pits. Therefore, in these traces clathrin intensity exceeds the detection threshold earlier than the P/S ratio, and converges into a plateau at earlier stages of the coat lifetime as the apex of the coat moves further away from the interface. Since the entire adherent surface of the plasma membrane is essentially parallel to the substrate, the background noise in the S channel overwhelms the signal (supplementary figure 2e in^33^) and increases the detection threshold of the P/S signal. Therefore, we propose that using P signal only (instead of P/S) would serve as a more accurate marker for determining the timing of curvature generation by endocytic clathrin coats.

Flat-to-curved transition and constant curvature models are proposed to explain formation of individual clathrin pits that are independent of other clathrin-coated structures. We therefore do not extend our conclusions to endocytic clathrin pits detected at the peripheral regions of plaques ^16,41^. Indeed, clathrin pits that appear to disassociate from larger clathrin assemblies are observable in TIRF-SIM acquisitions. However, in these assays, it is not possible to ascertain whether these structures are independent clathrin pits that happen to originate in the vicinity of a plaque ^15^.

It is important to note that clathrin plaques are predominantly observed at the adherent surfaces of cells and their abundance is dependent on adhesive properties of the substrate on which cells are cultured ^43^. In particular, polylysine treatment of coverslips, often used prior to correlative light and electron microscopy assays^19^, increases prevalence of clathrin plaques^5^. It is likely that, rather than being precursors of endocytic pits, the primary function of plaques is related to cell-substrate adhesion^6,16^ and subsequent mechanotransduction^13^. However, endocytic models based on electron micrographs often times omit this possibility^4,19^. Overall, our results signify the importance of employing methodologies comprising high resolution in both spatial and temporal dimensions for constructing dynamic models.

## Acknowledgements

We thank the Advanced Imaging Center (AIC) at Janelia Research Campus for access to their TIRF-SIM system. We are particularly indebted to Aaron Taylor and Satya Khuon from the AIC team. The AIC is jointly supported by the Howard Hughes Medical Institute and the Gordon and Betty Moore Foundation.

## Authorship

C.K., E.C., N.M.W. and J.P.F. conceived the study. C.K. performed the TIRF-SIM experiments on Drosophila embryos and SUM159 cells. E.B. and D.L. designed and performed high-NA TIRF-SIM measurements. J.P.F. developed the software for manual particle tracking and image analysis. J.P.F., C.C. and F.H. performed manual particle tracking and image analysis. N.M.W. has developed and implemented the automated clathrin trace analysis. H.C.C. developed UAS-mEmerald-Clc transgenic flies. A.T., S.L. and R.Z. have developed the self assembly model used in simulation. M.W. and Y.C. designed and performed single molecule imaging and FRAP assays on cell-free reconstitution systems. C.K., N.M.W. and J.P.F. wrote the manuscript.

## Data and materials availability

Datasets and software used in this study are available to the scientific community upon request to C.K.

## Materials and Methods

### Cell Culture

BSC-1 cells stably expressing EGFP-CLTA and COS-7 cells transiently expressing mEmerald-CLTB were cultured and prepared for microscopy as described in [1]. SUM-159 cells genome edited to express σ2-EGFP [2] and transiently expressing mRuby-CLTB (Addgene; Plasmid #55852) were cultured in F-12 medium with hydrocortisone, penicillin-streptomycin and 5% fetal bovine serum (FBS). Gene Pulser Xcell electroporation system (Bio-Rad Laboratories, CA, USA) was used for transient transfection of SUM-159 cells following manufacturer’s instructions. SUM-159 cells were cultured on 35mm glass bottom dishes (MatTek) and imaged 24-48 hours after transfection in phenol-red-free L15 (Thermo Fisher Sci.) supplemented with 5% FBS at 37°C ambient temperature.

### Drosophila Embryos

To construct pUAST-mEmerald-Clc, the in-frame GFP fusion from pUAST-GFP-Clc [3] was excised with an EcoRI/BglII digest and replaced with mEmerald (Addgene), PCR amplified with primers GGAATTCCACCATGGTGAGCAAGGGCGAGG and CGAGATCTGAGTCCGGACTTGTACAGCTCGTCCATG (the EcoRI and BglII sites in the primers are underlined). After verifying the resulting plasmid by sequencing (Purdue Genomics Core Facility), UAS-mEmerald-Clc transgenic flies were generated by P element-mediated transformation [4].

UAS-CLC-mEmerald flies were mated with a GAL4-Arm (Bloomington Drosophila Stock Center) line to produce embryos expressing CLC-mEmerald in the amnioserosa and surrounding germ-band tissue. The embryos were collected and aged for 10-12 hours at 25°C. After dechorionation, embryos were mounted on coverslips and immersed in voltalef oil for imaging at 22°C.

### Microscopy

High-NA TIRF-SIM images of COS-7 and BSC-1 cells were acquired with 1.7-NA 100x objective (Olympus Life Science, CA, USA) as described in [1]. SUM-159 cells and Drosophila embryos were imaged using the TIRF-SIM system at the Advanced Imaging Center (AIC) of the Janelia Research Campus. This system comprises a 1.49-NA 100x objective lens (Olympus Life Science, CA, USA) fitted on an inverted microscope (Axio Observer; ZEISS) equipped with a sCMOS camera (ORCA-Flash4.0; Hamamatsu). Structured illumination was provided by a spatial light modulator as described in [1] and movies were acquired using 20 ms exposure time and with frame rates ranging from 0.15 to 0.5 Hz. To image clathrin activity in tissues of Drosophila embryos, the apical surface of amnioserosa and germ-band cells were positioned within the evanescent illumination by gently pressing the dechorionated embryos against the coverslip.

Single triskelion recruitments were performed on cell-free clathrin assembly assays and detected using TIRF imaging as detailed in [5].

### Image Analysis

#### a. Manual analysis

Clathrin-coated structures in the TIRF (diffraction-limited) channel were tracked using two-dimensional cmeAnalysis software [6]. The traces extracted this way were filtered based on their intensity profiles as described in [7]. Briefly, to distinguish complete clathrin coat formation events, only the traces that contain both positive and negative intensity slopes (corresponding to coat growth and dissolution, respectively) were selected.

To analyze the images in the TIRF-SIM (superresolved) channel, a program was written in MATLAB 2017b (Mathworks) for the user to manually locate acceptable traces based on the following rejection criteria: 1) The structure must not come into contact with any other structure at any point during its lifetime (to exclude pair spots). 2) The structure must perform a single phase of intensity increase followed by a single phase of intensity decrease (to exclude hotspots). Once a structure was selected, the user defined the first and last frames of the structure’s existence. The *x-y* position of the structure was taken as the intensity-weighted center of the image. The area was calculated using the MATLAB built-in function, edge, set to calculate the Canny edge of a small window containing only the structure of interest. When the edge finder failed to close the edge, the algorithm allowed the user to close the edge manually (Fig. S7). The area within the closed boundary is calculated as the sum of the pixels enclosed.

Time averages of clathrin-coated pit traces are determined using TIRF-SIM images centered on the intensity-weighted center of the structure at each time point. These image sequences are of varying lengths, so each is extended to a length of 100 frames by linearly interpolating respective pixels at each time point (Fig. S8). Images corresponding to each time/completion point are averaged pixel-by-pixel to obtain the time averages shown in Movies 4 & 5. In each frame, the area is calculated using the Canny edge-detection algorithm as described above.

To get the radial averages shown in Figure 3, the images were upscaled by a factor of 10 with bicubic interpolation. For each upscaled image, 36 radial kymographs (separated by 5° increments and concentric with the intensity-weighted center of the structure; Fig. 9A) are averaged and fit to the sum of two Gaussian functions (Fig. S9B):

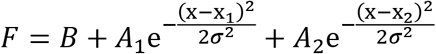

where *B* is the background, *A*_1_ and *A*_2_ are the amplitudes, *x*_1_ and *x*_2_ are the center points, and *σ* is the standard deviation. *σ* values for the largest area deciles are 40.6 nm for COS-7 Clathrin-mEmerald, 37.3 nm for BSC-1 Clathrin-EGFP, 68.7 nm for SUM-159 Clathrin-Ruby, 45.4 nm for SUM-159 AP2-EGFP and 51 nm for embryo Clathrin-mEmerald. As expected, the standard deviations are the smallest for the data sets obtained by high-NA TIRF-SIM (i.e. COS-7 and BSC-1) and the largest for the SUM-159 Clathrin-Ruby dataset due to longer fluorescence wavelength.

#### b. Automated analysis

##### Particle tracking

We used TraCKer [8] for automated tracking of clathrin-coated structures in TIRF-SIM (superresolved) channel. Since TraCKer is originally developed to trace diffraction-limited spots in fluorescence movies, curved clathrin structures that appeared as rings were frequently not detected. To circumvent this, as an initial step, holes (in rings) were closed using an averaging convolution followed by an erosion. After obtaining the traces of clathrin coats (*x-y* coordinated in multiple frames) all further analysis utilized the unmodified TIRF-SIM images.

##### Detection of ring patterns in TIRF-SIM Images

After tracking, each clathrin trace is evaluated for appearance of ring pattern using an 11×11 pixel window (~361nm × 361nm) centered on the structure in each frame (Fig. S10A & C). A coordinate pair (r_i_, Int_i_) is defined for each pixel in the cropped image, where r_i_ is the distance from the center of the i^th^ pixel to the center of the structure and Int_i_ is the intensity of that pixel. Plotting these coordinates gives a radial profile of the structure to which a Gaussian distribution is fit using least squares (Fig. S10B & D). If the peak of the Gaussian is determined to be at least 2 pixels (60 nm) from the center of the structure, the structure is considered to have the ring pattern (Fig. S10B). Only the traces that have the ring pattern in two consecutive frames and no ring pattern in the first frame are used in further analysis. The latter constraint is applied because of the in analyzing the frames leading up to maximum curvature.

##### Boundary detection

The average image cross-section is determined by averaging 36, equidistant, radial crossections (Fig. S11A & F). This 1D profile image was fit to the sum of two Gaussians (Fig. S11D & I), and the boundary of the structure was determined by the points where the magnitude of the Gaussians’ slopes were found to be a maximum (Fig. S11E & J).

#### Self-assembly model

Figure S12 shows the schematic representation of clathrin triskelions used in simulations, where *D*_*hh*_ (~18.5 nm) is the nearest neighbor hub-hub distance (i.e., “strut” length) and 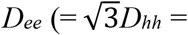 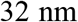; ***ϕ*** ≈ 120 is the next to nearest hub-hub distance [9, 10]. From basic topology, any closed shell will contain 12 pentagons. If we further assume that those pentagons arrange so that there is icosahedral symmetry, then all possible configurations are classified by the *T* number according to

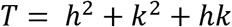

with *h* and *k* non-zero positive integers. Then, the clathrin coat consists of 20*T* triskelions, 30*T* “struts” of length *D*_*hh*_ and 10*T*+2 polygonal faces; 10(*T*-1) of them hexagons and 12 pentagons. The radius *R* of the clathrin coats will therefore be defined by the distance between the hub and the center of the circumscribed sphere, which is given by

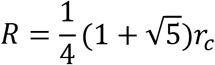

where *r*_*c*_ is the distance between two nearest neighbor pentagons. This distance may be evaluated as _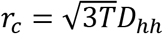_ and therefore, we have a relation between the sphere radius *R* and *D*_*hh*_ according to

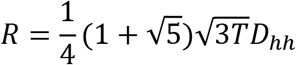

We adapt a previous model that successfully described the self-assembly of capsid subunits as assisted by scaffolding proteins [11]. Here, the role played by the latter are the AP2 adapters and the former are clathrin triskelions. The model is defined by

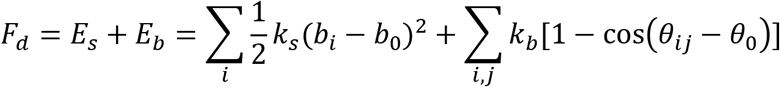

with *θ*_0_ a preferred angle, related to the spontaneous curvature *H*_0_. The stretching energy sums over all bonds *i* with *b*_0_ the equilibrium bond length and the bending energy is between all neighboring triskelions indexed with *i*. We further assume that there is an attractive force between the triskelions and the preformed scaffolding layer (AP2 subunits), which involves a simple LJ-potential 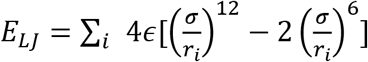 with *ϵ* the depth of the potential and *σ* the position of minimum energy corresponding to optimal distance between the center of the core and the subunits. The mechanical properties of the subunits (the ratio of stretching to bending moduli) are described by the Foppl von-Karman (FvK) number that is defined by

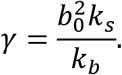

In our simulations (Figs. 1B&D, Movies S1&2), FvK number *γ* =100 is selected based on the previous work [11]. It is important to remark that the pentagons are dynamically generated as the simulation proceeds and therefore, their location is an unambiguous prediction by the geometry and theory (Fig. 1F). It should be emphasized that, in contrast with the situations for viruses [12], clathrin shells may not exhibit icosahedral order. Still, the mechanism of defect growth is *universal* (independent of FvK number) and the absence of icosahedral order does not qualitatively modify any of the conclusions.

Movies displaying the geometry of clathrin coat assembly (Movie S1&S2) were rendered in MATLAB using an orthographic projection. In the case of curved structures, the relative positions of clathrin triskelia are given by the results of the self assembly model. After the addition of each new triskelion the entire structure is rotated so that the position vector of the center of mass moves towards vertical. For flat structures, triskelia are added in any position in the hexagonal lattice such that the distance from the new triskelion to the center of mass is minimized. When new triskelia are added, the structure translates in order to move the center of mass towards the origin. The strut length is consistent in all structures.

#### TIRF-SIM simulations

High-NA TIRF-SIM simulations in Figures 1 & S5 and Movies S1 & S2 were created by modeling the fluorescence of each labeled triskelion as a 3D Gaussian. The amplitude of the each Gaussian is proportional to the intensity of the incident light at its position within evanescent TIR field, in which the intensity profile used is an exponential decay along the z-axis. The image is generated by calculating the sum of all emitted intensities at the central position of each image pixel at the focal plane. All numeric parameters were implemented as reported in [1]: TIRF decay length = 50 nm; XY Gaussian SD = 84 nm; Z Gaussian SD = 500 nm; image pixel size = 30 nm.

#### Clathrin remodeling simulations

In every step of the simulations shown in Figure 7B, clathrin triskelia are stochastically removed and then added to the coat. The probability of removal and addition at a particular location is based on the number of triskelia currently in the coat neighboring that location. For the simulation where remodeling is limited to the rim of the coat, the probabilities of removal used are 0, 1/23, 1/23, and 0 corresponding to the location having 0, 1, 2, or 3 neighboring triskelia respectively. Similarly the probabilities of removal in the simulation where remodeling can occur anywhere are 0, 1/120, 1/120, 1/120. Probabilities of addition for both simulations are 0, 0.1, 0.1, 0.1. The geometry of the coat being constructed is a T=7 incosahedrally symmetric polyhedron. When if, at the end of a simulation step, the coat is found to be complete (every vertex of the polyhedron is occupied), the simulation ends.

---

**Figure S1.**
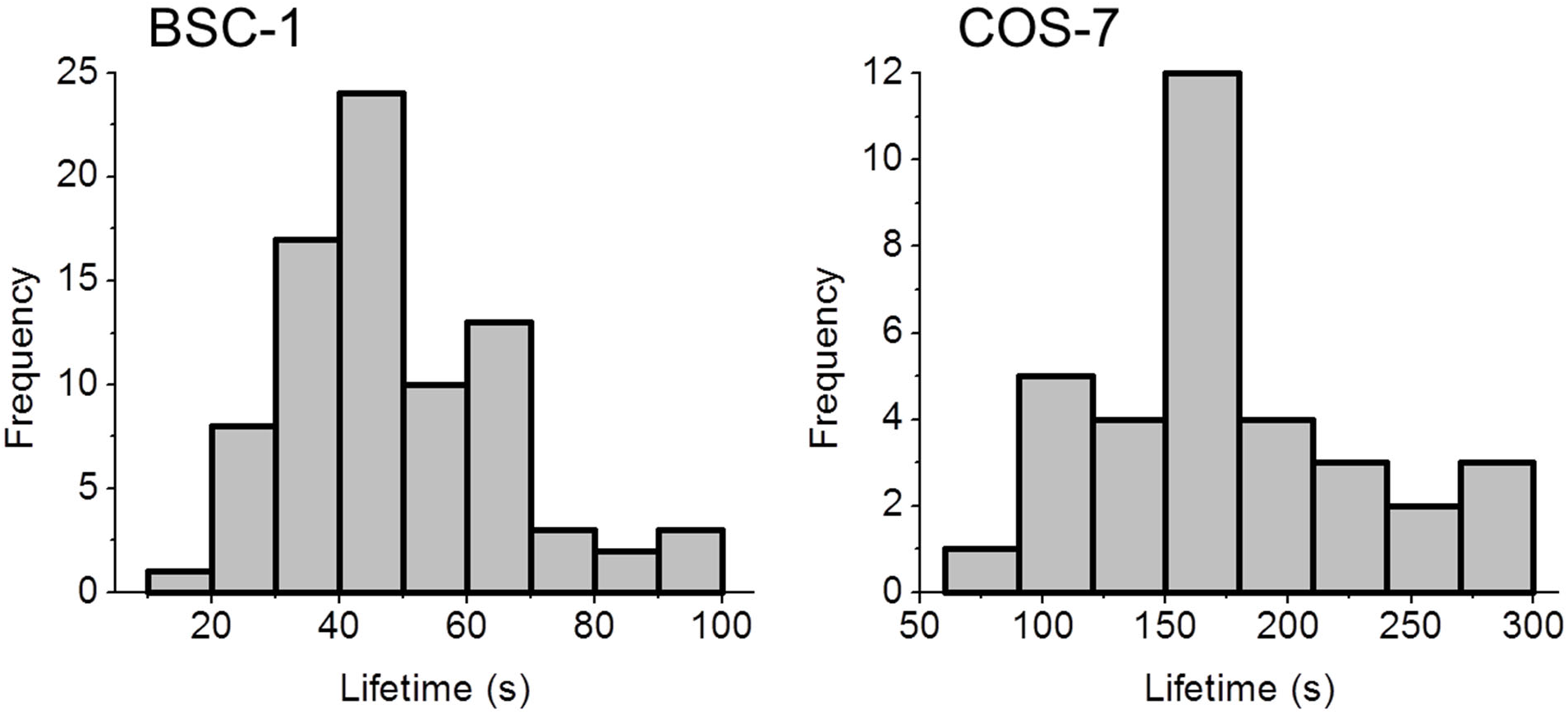
Lifetime distributions of manually selected clathrin-coated pit traces extracted from high-NA TIRF-SIM acquisitions of BSC-1 cells expressing clathrin-EGFP (left) and COS-7 cells expressing clathrin-mEmerald (right).

**Figure S2.**
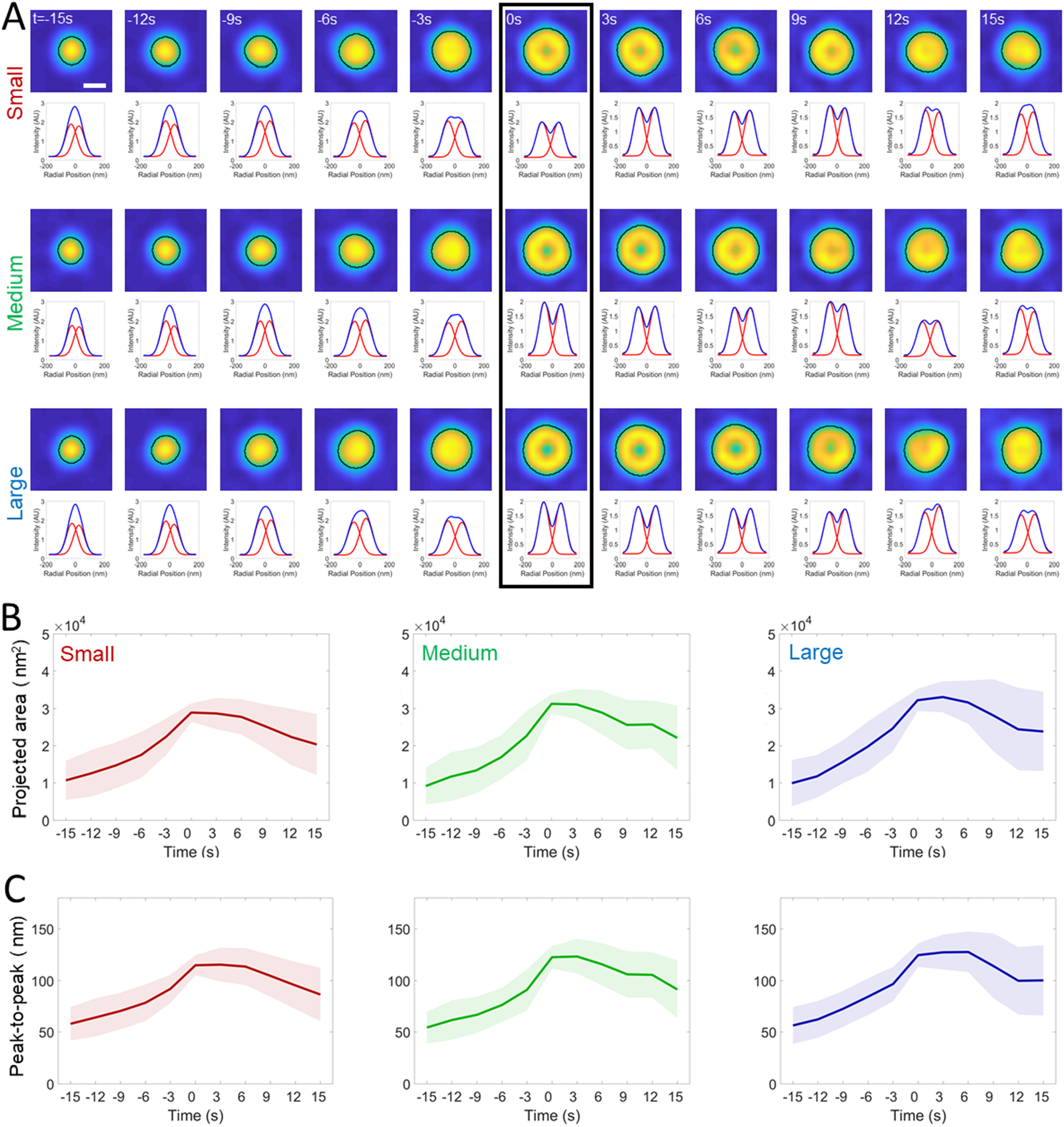
Automated analysis of the projected area during formation of clathrin pits in BSC-1 cells. **A.** 210 traces detected from two BSC-1 cells (imaged at 0.33 frames/second) are grouped and synchronized using the first detected ring pattern (black rectangle). Average images (top) and corresponding radial averages (bottom) are shown for each group (small, medium and large). Detected boundaries are shown by the black demarcations. The temporal evolution of the average area **(B)** and peak-to-peak separation of the Gaussian fits to the radial averages **(C)** are plotted for the three groups. Shaded areas represent the standard deviation. Scale bar, 100 nm.

**Figure S3.**
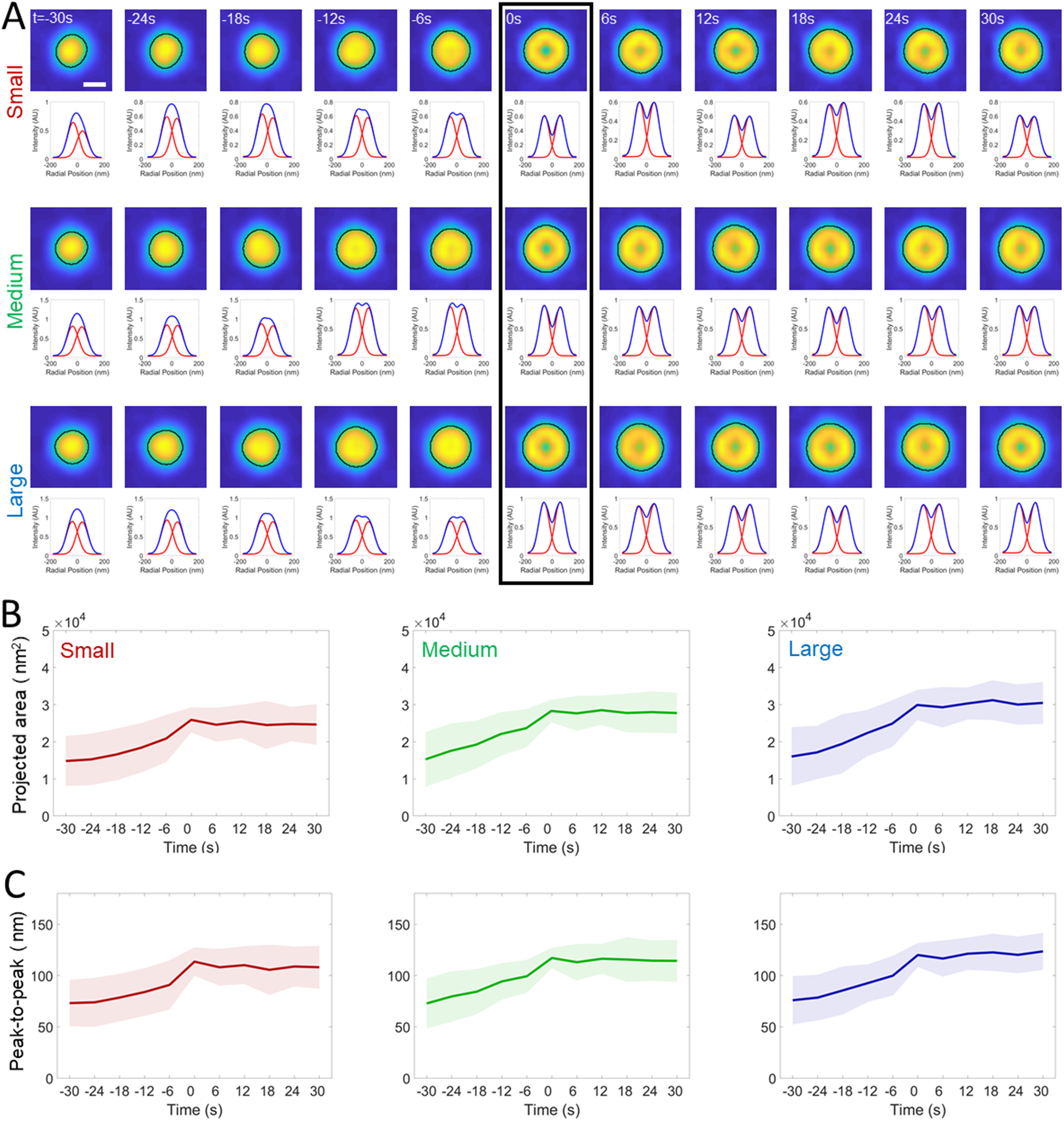
Automated analysis of the projected area during formation of clathrin pits in COS-7 cells. **A.** 519 traces detected from one COS-7 cell (imaged at 0.167 frames/second) are grouped and synchronized using the first detected ring pattern (black rectangle). Average images (top) and corresponding radial averages (bottom) are shown for each group (small, medium and large). Detected boundaries are shown by the black demarcations. The temporal evolution of the average area **(B)** and peak-to-peak separation of the Gaussian fits to the radial averages **(C)** are plotted for the three groups. Shaded areas represent the standard deviation. Scale bar, 100 nm.

**Figure S4.**
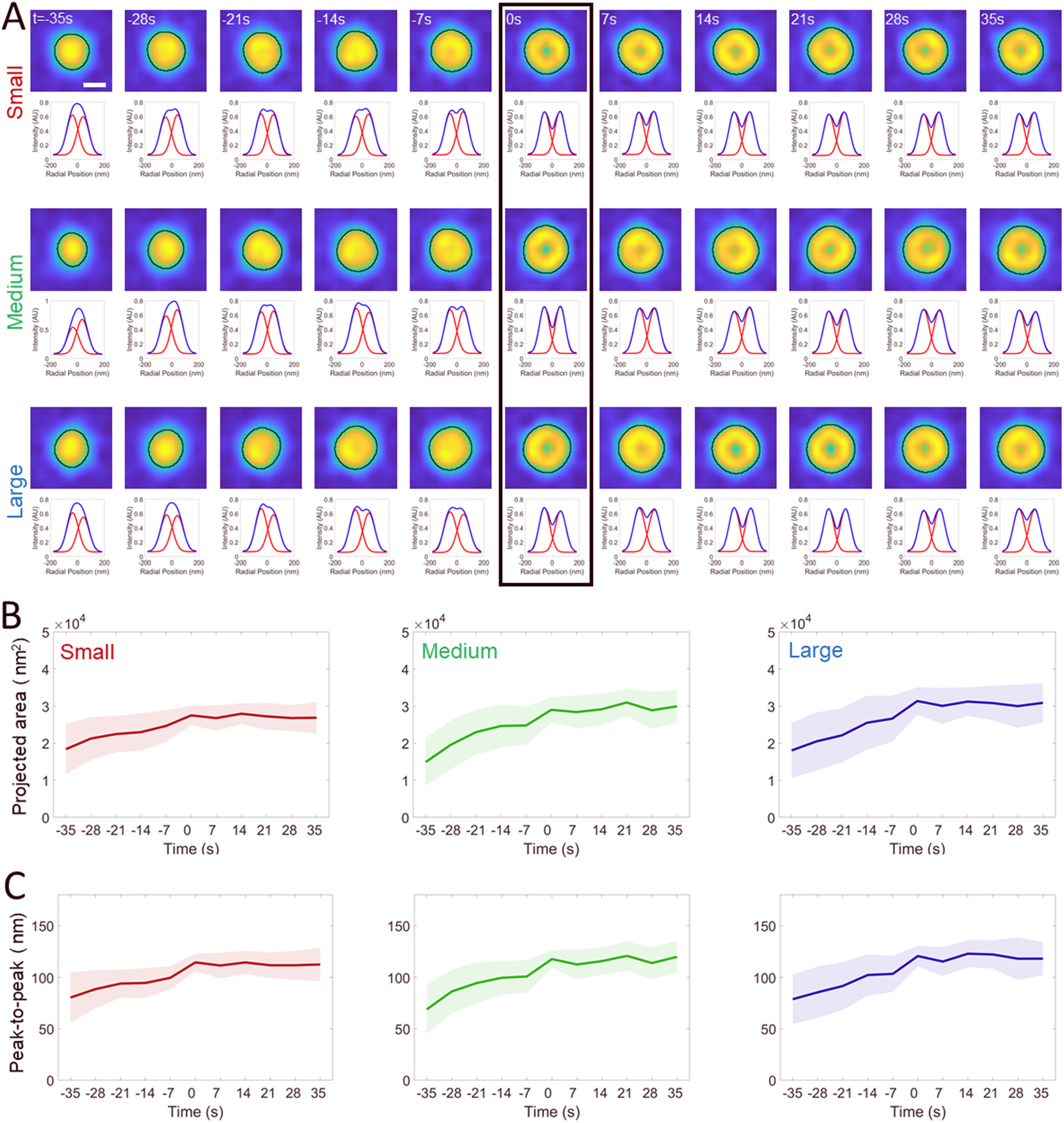
Automated analysis of the projected area during formation of clathrin pits in COS-7 cells. **A.** 175 traces detected from one COS-7 cell (imaged at 0.2 frames/second) are grouped and synchronized using the first detected ring pattern (black rectangle). Average images (top) and corresponding radial averages (bottom) are shown for each group (small, medium and large). Detected boundaries are shown by the black demarcations. The temporal evolution of the average area **(B)** and peak-to-peak separation of the Gaussian fits to the radial averages **(C)** are plotted for the three groups. Shaded areas represent the standard deviation. Scale bar, 100 nm.

**Figure S5.**
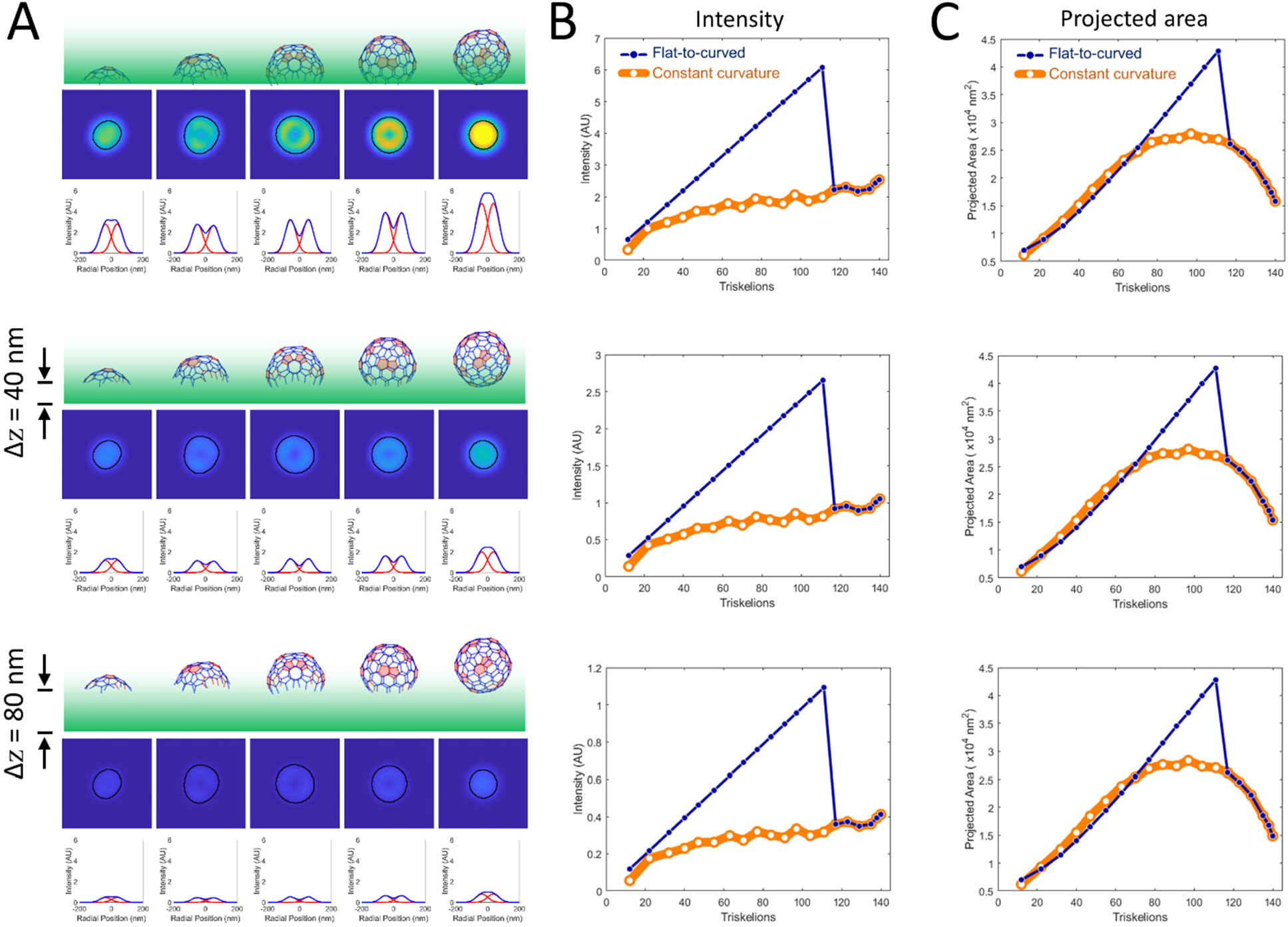
Flat-to-curved transition results in a substantial decrease in the TIRF signal. **A.** High NA TIRF-SIM simulations and corresponding radial averages are shown for different stages of T=7 polyhedron formation according to self-assembly model. Simulations are performed using 50 nm penetration depth of the evanescent field (green), and offsets of 0 (upper), 40 (middle) and 80 nms (lower) between the interface and the clathrin coat. **B.** Corresponding intensities are plotted for the simulations in A. Regardless of the distance between the interface and the coat, the constant curvature predicts a plateau in the clathrin signal (orange), whereas a late stage flat-to-curved transition (at 70% of completion) results in a significant decrease in intensity (blue). **C.** Area within the detected boundaries are plotted with respect to triskelion numbers for the corresponding simulations in A. Regardless of the distance between the substrate and the coat, the constant curvature predicts a plateau in the detected area (orange), whereas a late stage flat-to-curved transition (at 70% of completion) results in a significant decrease in area (blue).

**Figure S6.**
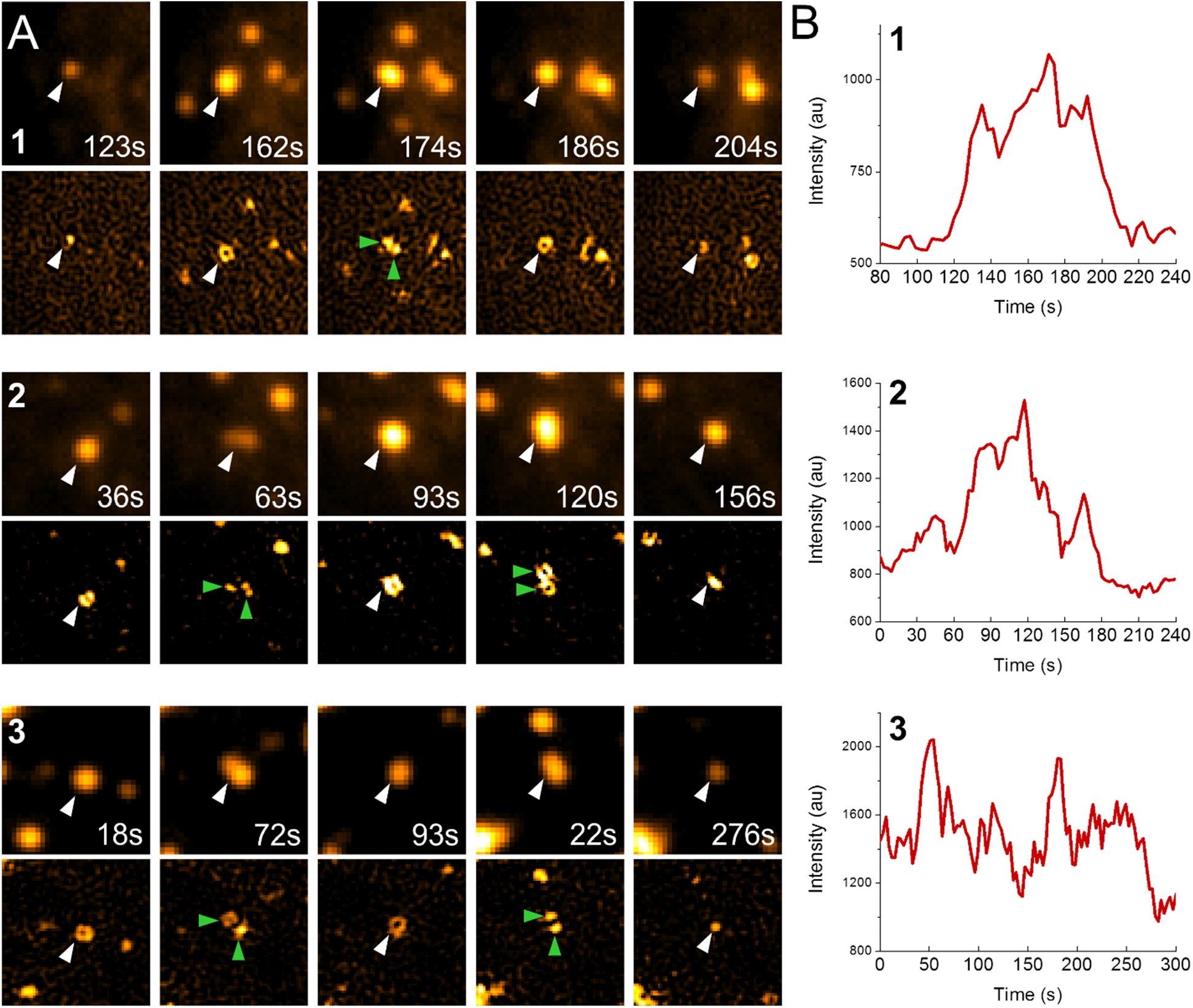
Fluorescent spots that appear as individual structures under diffraction-limited imaging may correspond to multiple independent clathrin-coated pits. **A.** Three examples of diffraction-limited fluorescent spots comprising multiple clathrin-coated structures. Top rows show TIRF snapshots of individual diffraction-limited spots (marked by white arrowheads) at different time points. Bottom rows correspond to high-NA TIRF-SIM images of the same region, where green arrowheads mark multiple structures in close proximity that cannot be resolved in the TIRF channel (top row). **B.** Plots show the fluorescence intensity traces of the diffraction-limited spots in A.

**Figure S7.**
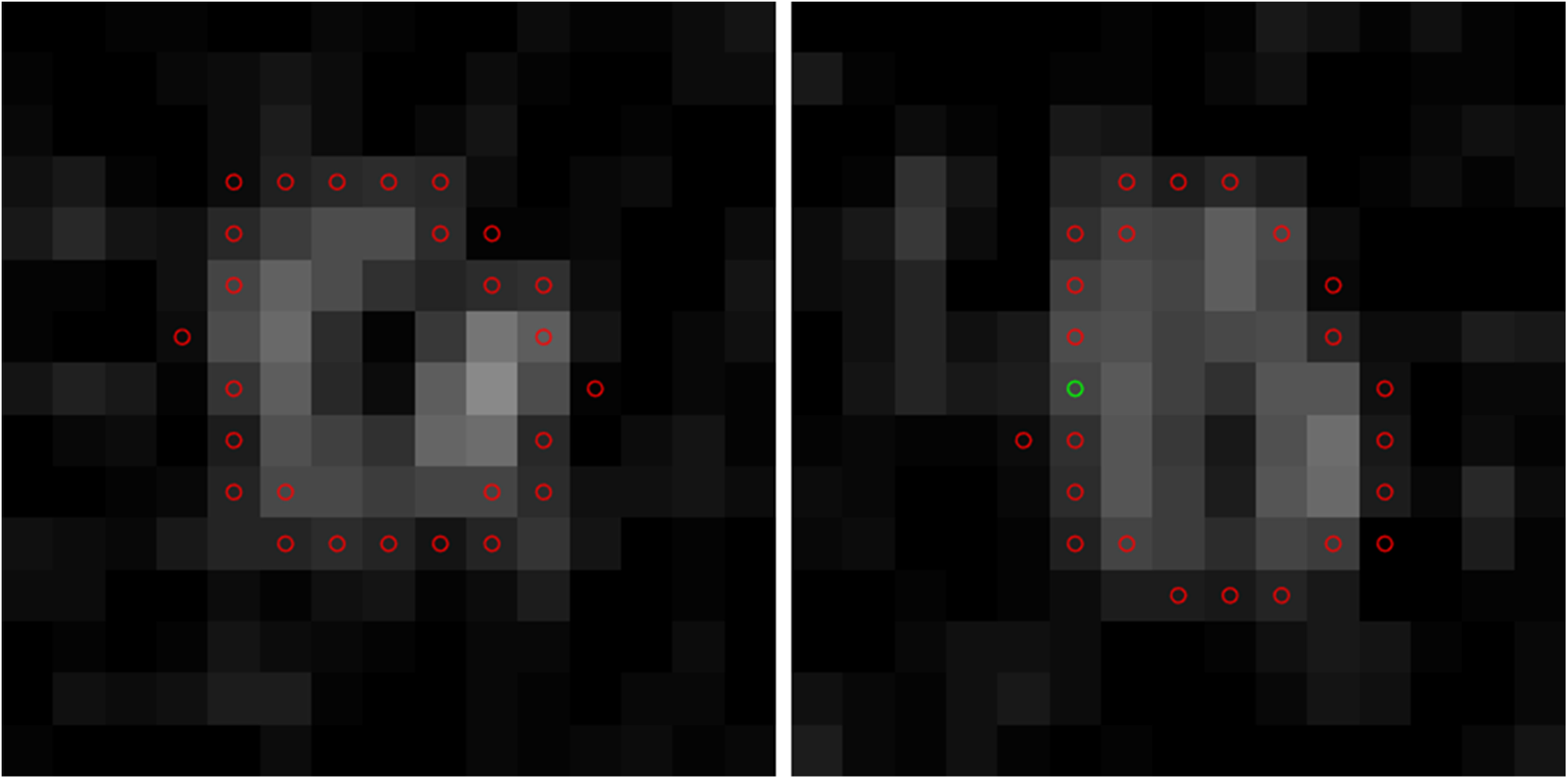
Red circles in each image correspond to the edges detected by the Canny edge-detection algorithm. The pixels within this boundary are used to define the designated area of the structure (left). When the Canny algorithm fails to close the boundary (right), the user has the option to manually assign the missing edge pixels (green circle).

**Figure S8.**
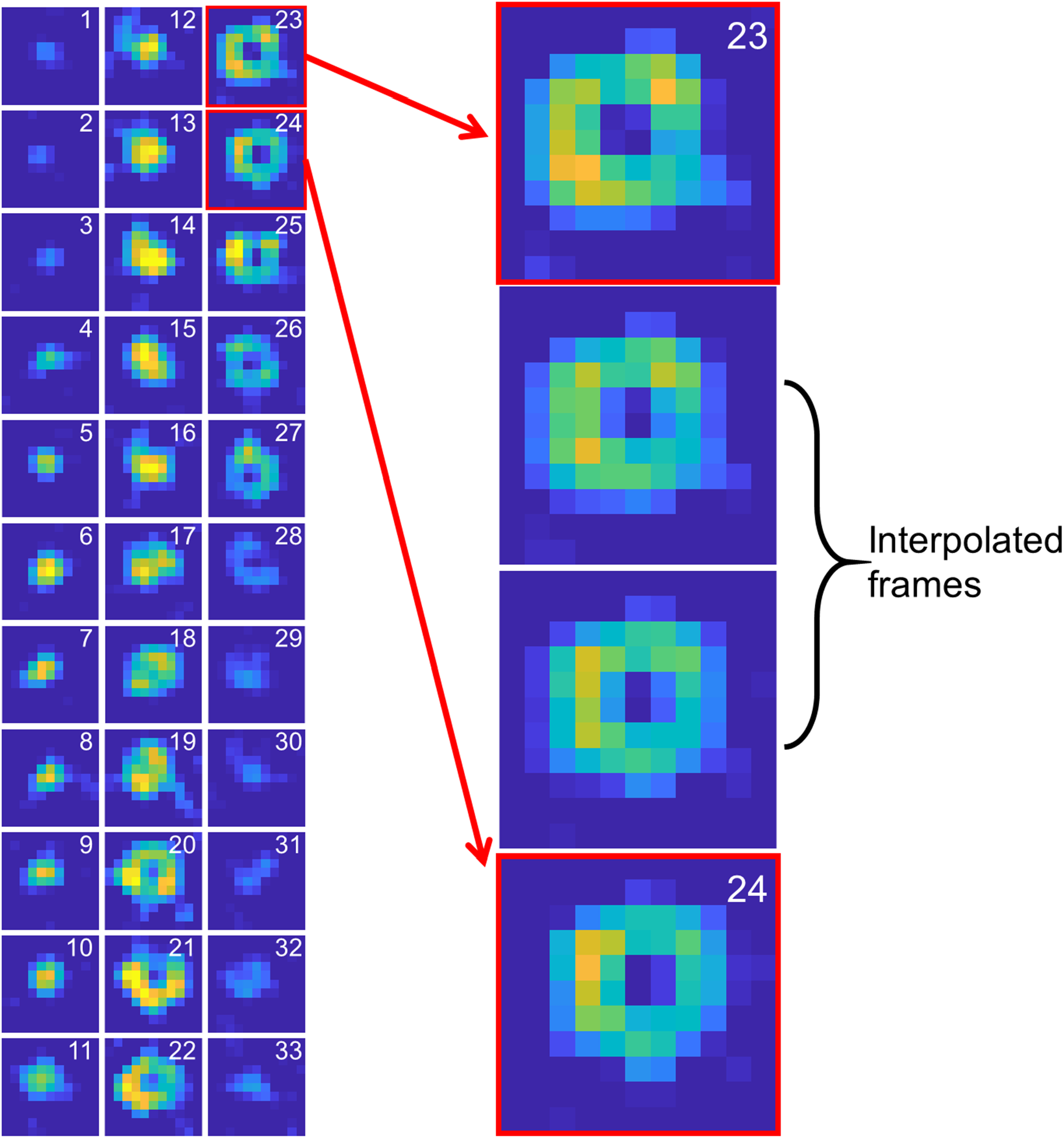
33 consecutive TIRF-SIM images corresponding to formation and dissolution of a clathrin-coated pit in a COS-7 cell expressing clathrin-mEmerald (left). Blowup image shows the linear interpolation linking frames number 23 and 24 (right).

**Figure S9.**
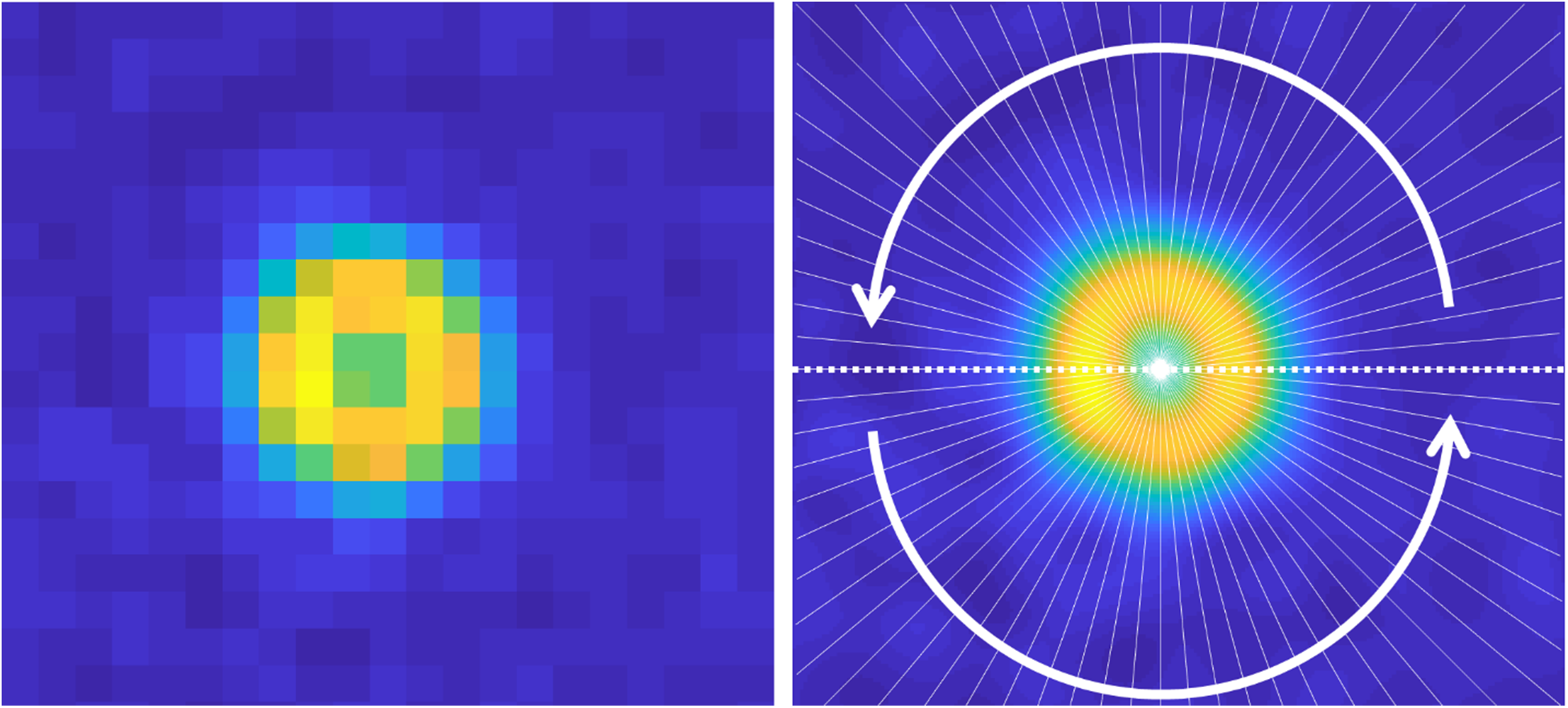
An average clathrin-coated pit image (left) and its upscaled version (right). Kymographs are calculated along the radial white lines separated by 5°. Radial averages are determined for different conditions by averaging the radial kymographs.

**Figure S10.**
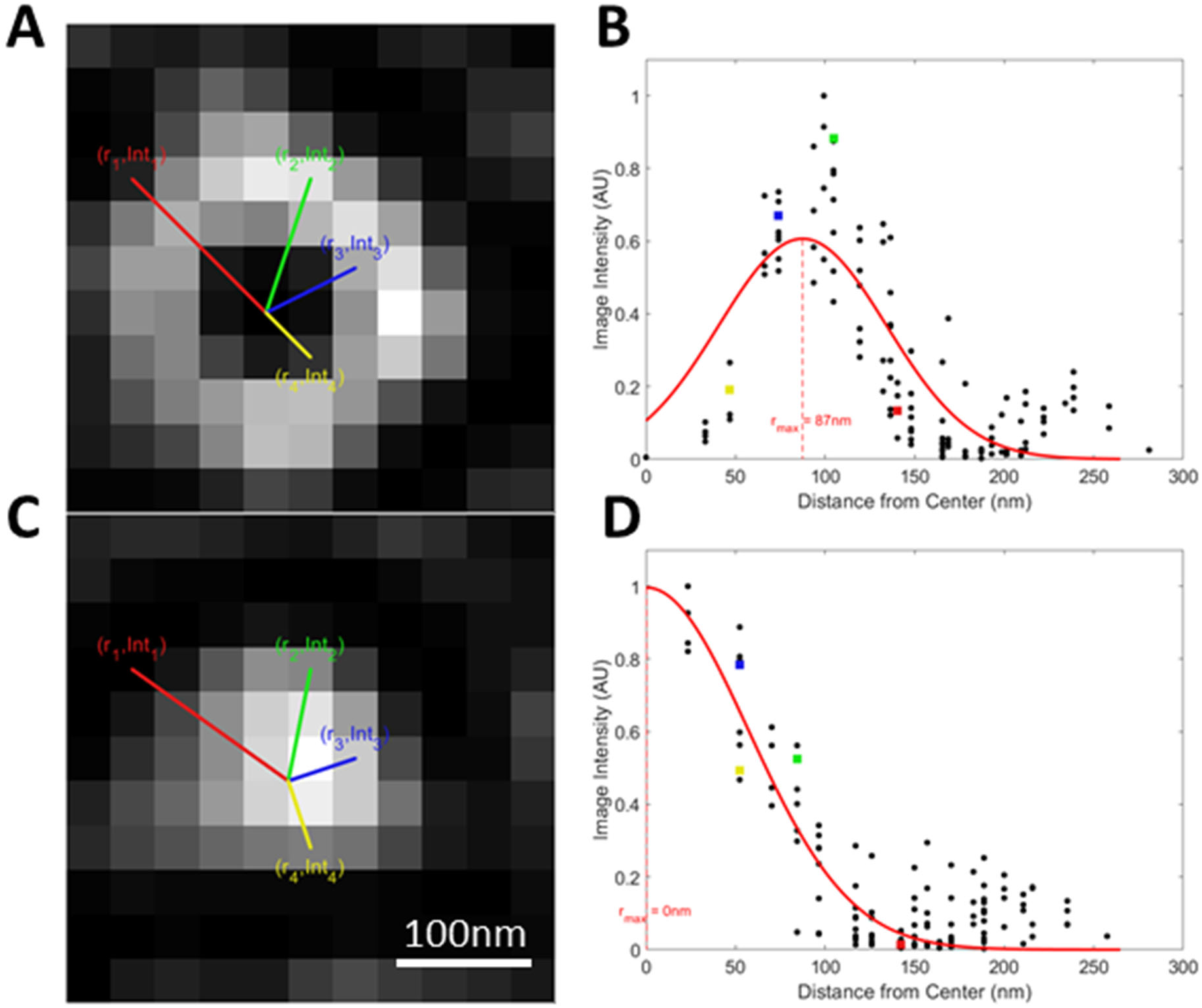
**A & C.** For each time point in the trace, the analysis is performed within the 11×11 pixel region around the center of the structure. **B & D.** A coordinate pair (r_i_, Int_i_) is defined for each pixel in the cropped image, where r_i_ is the distance from the center of the i^th^ pixel to the center of the structure and Int_i_ is the intensity of that pixel. Plotting these coordinates gives a radial profile of the structure. (Colored squares in B, D are the same colored pixels indicated in A, C respectively). To this profile, a Gaussian distribution is fit using least squares. The mean value of this Gaussian is the radial distance of maximum intensity (r_max_). When this r_max_ value is above a threshold (empirically chosen to be 60 nm), we can be confident that the structure displays obvious curvature (**B**). Below this threshold we find that curvature cannot be resolved (**D**).

**Figure S11.**
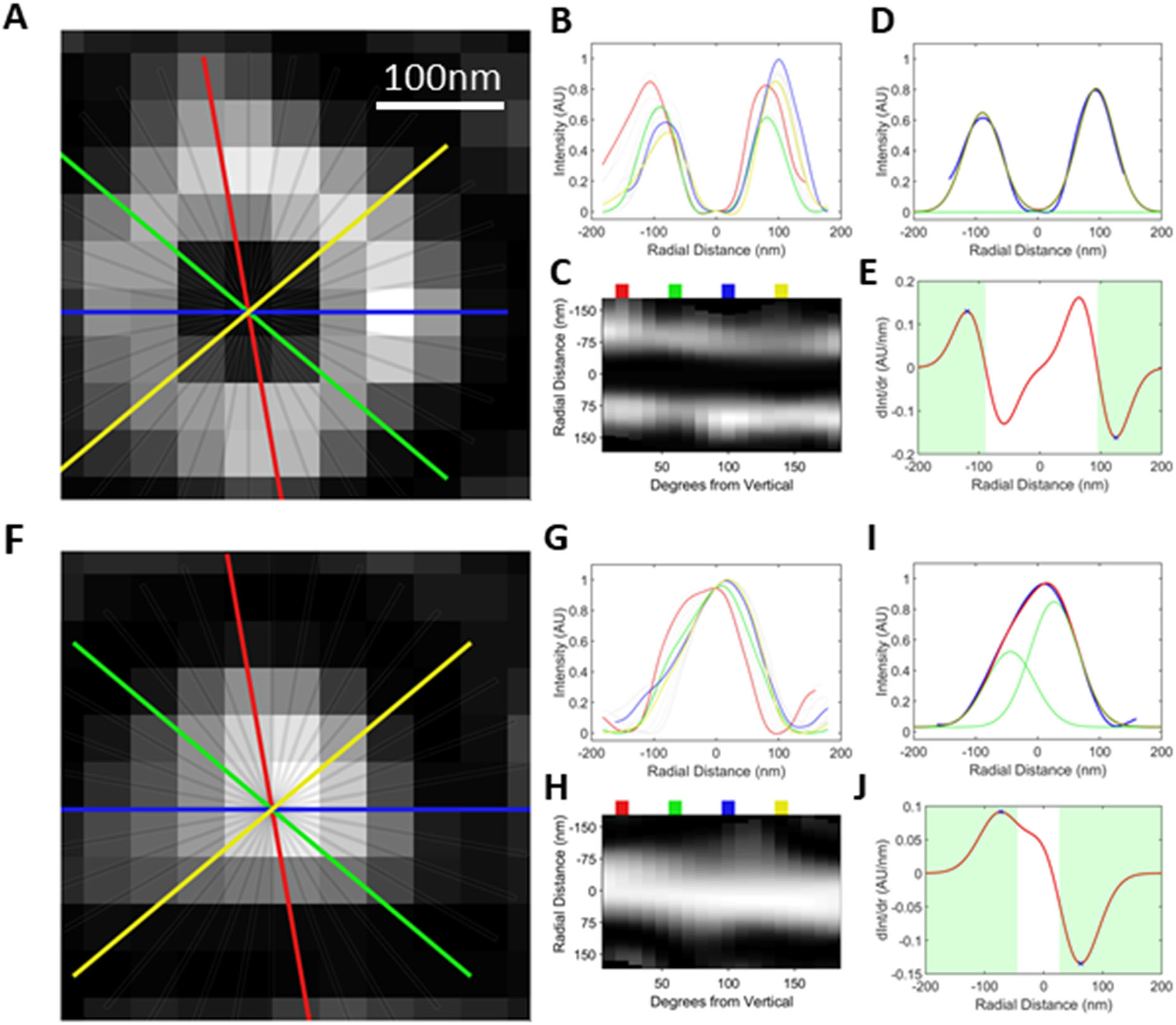
For boundary detection, we calculate the profiles of 36 lines running through the center of the clathrin-coated structure (only 18 lines are shown in **A** and **F**) in each time point. The profiles can be viewed as line plots (**B, G**) or as images showing the profile intensity on one axis and the angle of the profile line (from vertical) on the other (**C, H**). The line plots (**B, G**) and the angles marked with a colored boxes (**C, H**) are the representations of the corresponding colored profiles (**A, F**). To calculate the area of the clathrin coat, we take the average of these profiles and fit it to the sum of two Gaussian distributions (which are required to have the same standard deviation) using least squares (**D, I**; blue is the average profile; red is the fitted Gaussians; green are the fitted Gaussian separated). The average extent of the clathrin coat is chosen to be the maximum and minimum slopes of the fit. We take the derivative of the fit (**E, J**; red line). In order avoid finding a slope which is on the interior of the profile, we consider slopes which exterior to the Gaussian peaks (**E, J**; shaded green regions). The maximum slope in the left region and the minimum slope in the right region define the average diameter of the clathrin coat (**E, J**; blue crosses).

**Figure S12.**
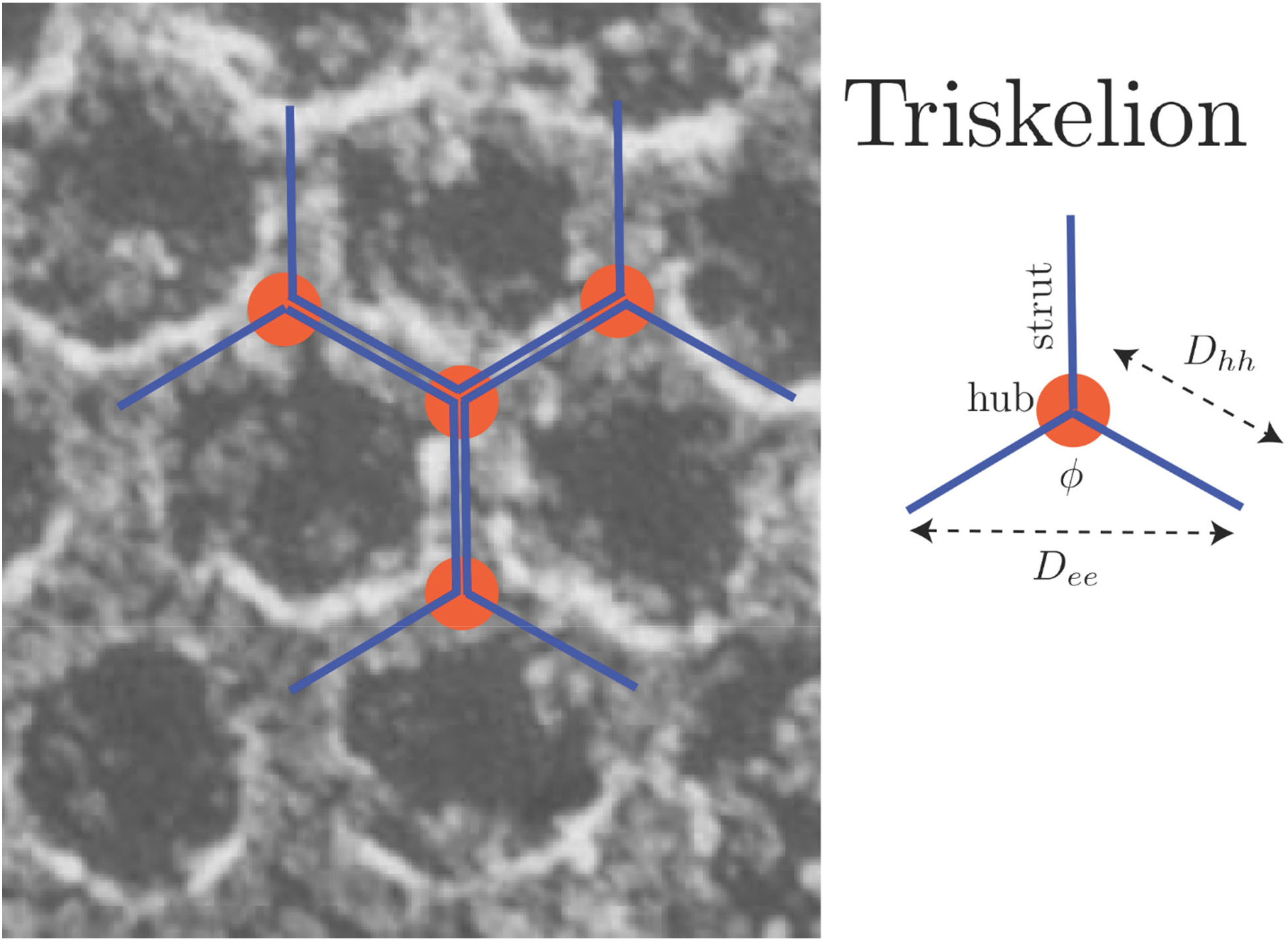
Schematic representation of a clathrin triskelion superimposed to an electron micrograph [8].

**Movie S1.** Assembly of a T=7 polyhedron according to the constant curvature model, where the position of each new triskelion is determined using the self assembly model. The coat growth is shown in the *x-z* and *x-y* planes in **A** and **B**, respectively. In **A**, the color bar represents normalized evanescent TIR field intensity. Fully assembled pentagons are highlighted in pink. **C.** Simulated high-NA TIRF-SIM images corresponding to the clathrin coats shown in A and B. The red line shows the detected boundary. **D.** Projected area (i.e., area within the detected boundary) is plotted with respect to the number of triskelions integrated into the coat. Scale bars, 20 nm.

**Movie S2.** Assembly of a T=9 polyhedron according to the constant curvature model, where the position of each new triskelion is determined using the self assembly model. The coat growth is shown in the *x-z* and *x-y* planes in **A** and **B**, respectively. In **A**, the color bar represents normalized evanescent TIR field intensity. Fully assembled pentagons are highlighted with pink. **C.** Simulated high-NA TIRF-SIM images corresponding to the clathrin coats shown in A and B. The red line shows the detected boundary. **D.** Projected area (i.e., area within the detected boundary) is plotted with respect to the number of triskelions integrated into the coat. Scale bars, 20 nm.

**Movie S3.** TIRF-SIM movie of clathrin coat dynamics acquired at the amnioserosa tissue of a late Drosophila embryo expressing clathrin-mEmerald. Further zoom-in (blue box) shows growth and dissolution of an individual clathrin-coated pit (arrowhead).

**Movie S4.** Average clathrin-coated pit trace obtained from COS-7 cells expressing clathrin-mEmerald (left; N_traces_ = 34), where the black line is the detected boundary. The maximum projected area (area within the boundary) is reached at 75% of completion. For each image the radial average (red circles) is plotted on the right panel. Gaussian fits (green and blue) and corresponding peak-to-peak distances are shown between 40-96% completion.

**Movie S5.** Average clathrin-coated pit trace obtained from BSC-1 cells expressing clathrin-EGFP (left; N_traces_ = 81), where the black line is the detected boundary. The maximum projected area (area within the boundary) is reached at 69% of completion. For each image the radial average (red circles) is plotted on the right panel. Gaussian fits (green and blue) and corresponding peak-to-peak distances are shown between 40-96% completion.

**Movie S6.** TIRF-SIM (left) and TIRF (right) images of multiple clathrin-coated structures that appear as a single spot under diffraction-limited imaging. The movie corresponds to the second example in Figure S6.

